# Alternative probe chemistries for single-molecule analysis of long non-coding RNA

**DOI:** 10.64898/2025.12.04.691911

**Authors:** Kalika R. Pai, Aimee M. Martin, Madison Kadrmas, Julia R. Widom

## Abstract

Single-molecule microscopy has been widely used to study the structure and dynamics of RNA, but extension to larger systems such as long non-coding RNA (lncRNA) has proven challenging. Methods such as single-molecule kinetic analysis of RNA transient structure (SiM-KARTS), where the binding of a short, complementary oligonucleotide probe is used to determine accessibility of a specific region of the RNA, are promising. However, adapting SiM-KARTS to systems as complex as lncRNA requires careful optimization of experimental variables that have not been thoroughly explored. In this work, SiM-KARTS, thermal denaturation experiments, and circular dichroism spectroscopy were used to analyze the binding behaviors of probes with alternative backbone chemistries, specifically DNA with locked nucleic acid (LNA) residues incorporated and morpholinos. A segment of lncRNA that enabled control over the accessibility of the target sequence was used as a model. We show that optimizing probe backbone chemistry can allow for a more precise distinction between different structures of the target RNA, and for fine-tuning of probe binding stability without the structural impacts that other variables such as ionic concentration may have. Specifically, we demonstrate that LNA probes exhibit a high degree of structural sensitivity in both their binding and unbinding kinetics. We further show that when binding and unbinding rates are considered holistically, LNA probes allow traces arising from different target RNA structures to be individually classified with a high degree of accuracy. These results provide design principles for the application of SiM-KARTS to target RNAs of increased complexity such as lncRNA.

## Introduction

Long non-coding RNAs (lncRNAs) are RNAs that do not encode protein and are greater than 200 nucleotides in length. lncRNAs perform a variety of biological functions through interactions with DNA, RNA, or proteins. Specifically, many lncRNAs have regulatory functions, either epigenetic (1), transcriptional (2), or post-transcriptional (3). Structure is an important factor in mediating the interactions that allow for these regulatory functions (4). Additionally, many lncRNAs have been found to play roles in human disease, making the molecular details of lncRNAs interesting for the potential development of therapeutics (5–7). However, there have not been in depth studies of the structure or dynamics of most lncRNAs (8), which are difficult to study due to their large size and resulting structural complexity.

Single-molecule methods have strong potential for investigating the structure and dynamics of lncRNAs. Specifically, single-molecule microscopy avoids ensemble averaging and allows for the investigation of the dynamics of heterogeneous systems on time scales relevant to large structural rearrangements (9). However, common techniques, such as single-molecule Förster resonance energy transfer (smFRET) (10), are difficult to adapt to lncRNAs due to their length and structural complexity. For example, performing site-specific labeling of the RNA of interest can be costly, require complex experimental workflows, and impact the structures being studied.

An alternative to smFRET is single-molecule analysis of RNA transient structure (SiM-KARTS), which utilizes a short, fluorescently-labeled oligonucleotide probe that is complementary to a region of interest on the RNA (11–13). By tracking binding and dissociation of this probe, the accessibility of the region of interest can be revealed. For example, frequent observation of probe binding indicates high accessibility, implying a single-stranded local structure. SiM-KARTS can also be used to investigate conformational changes of RNA by evaluating variations in probe binding frequency. Using spike train analysis (14), “bursts” of probe binding can be distinguished from random events, uncovering transitions between conformations in which the target sequence is more or less accessible for probe binding. So far, SiM-KARTS has primarily been utilized to study riboswitches. For example, SiM-KARTS was performed on mRNA containing the preQ_1_ riboswitch using a DNA probe complementary to the Shine-Dalgarno (SD) sequence (termed an “anti-SD” probe) to monitor ligand-induced pseudoknot docking (11). It was determined that there are dynamic fluctuations in the accessibility of the target sequence, even at saturating ligand concentrations. In the case of the preQ_1_ riboswitch, techniques such as small angle x-ray scattering (SAXS), smFRET, and molecular dynamics simulations aided in experimental design (15, 16).

SiM-KARTS shows promise as a method for probing lncRNA as it removes the requirement for site-specific labeling. The RNA can instead be transcribed and then end-labeled using established chemistry performed under native conditions (17, 18), or a labeled capture oligonucleotide can be utilized to localize immobilized molecules without any modification to the target RNA. All previous reports of SiM-KARTS utilized an anti-SD probe, exploiting the presence of this motif in multiple riboswitches (19–21). For lncRNA, no such common motif exists, and multiple structural domains may be of interest. For this reason, to adapt SiM-KARTS for applications to lncRNA, designing an appropriate probe is of high importance. Challenges in probe design include ensuring binding is strong enough to yield clear optical signals, yet transient enough to pose minimal disruption to the target RNA’s dynamics. Lengthening the probe can increase binding stability, but also limits the precision with which specific regions of interest can be probed and increases the potential for the probe to disrupt the structure of the RNA. Mono- and divalent cations are well known to increase melting temperatures of DNA and RNA duplexes and hybrids (22), making adjusting cation concentrations a possible avenue for optimizing probe binding stability. However, as ionic conditions also have a large impact on stability of RNA secondary and tertiary structure (23), controlling probe binding stability in this manner could distort the structure of the RNA being studied.

One solution to fine-tune probe binding strength without changing the length, location, or structure of a target site is to consider alternative probe chemistries. Specifically, the incorporation of locked nucleic acid (LNA) residues into DNA is known to increase the melting temperatures of its hybrids with RNA and DNA (24). Incorporating one or two LNA residues into a short oligonucleotide can raise its predicted melting temperature by multiple degrees celsius, depending on placement within the strand (25). The binding strength of short sequences can also be increased by the use of morpholino oligonucleotides, which are typically used as gene knockdown reagents due to their high binding specificity and stability (26). In this study, we investigated the impact of backbone chemistry on the binding behavior of SiM-KARTS probes in different structural contexts and under different ionic conditions using a model RNA based on the lncRNA *Braveheart* (Bvht; Fig. 1A) (27–29). Bvht is a heart-specific lncRNA that was identified to be a regulator of cardiovascular commitment, functioning in the same regulatory pathway as the transcription factor *MesP1* during embryonic stem cell differentiation in mice (27). The structure of Bvht has been studied using chemical probing and SAXS, which revealed important features such as a 5’ asymmetric G-rich internal loop that interacts with a zinc-finger transcription factor called cellular nucleic acid binding protein (28). We show that probes with LNA residues incorporated demonstrate improved binding stability and structural sensitivity over DNA alone, while morpholino probes showed increased binding stability, but little structural sensitivity. We report a clustering approach in which the binding and unbinding rates observed within an individual trace are used as a kinetic fingerprint and show that these fingerprints effectively distinguish traces arising from different target RNA structures. Finally, we demonstrate that changes in the ionic environment can produce non-monotonic changes in probe binding kinetics, likely due to competing effects on the structure of the target RNA and the stability of the probe:RNA hybrid. These results will guide probe design for future studies utilizing SiM-KARTS, including investigations of lncRNA structure and dynamics.

**Figure 1.**
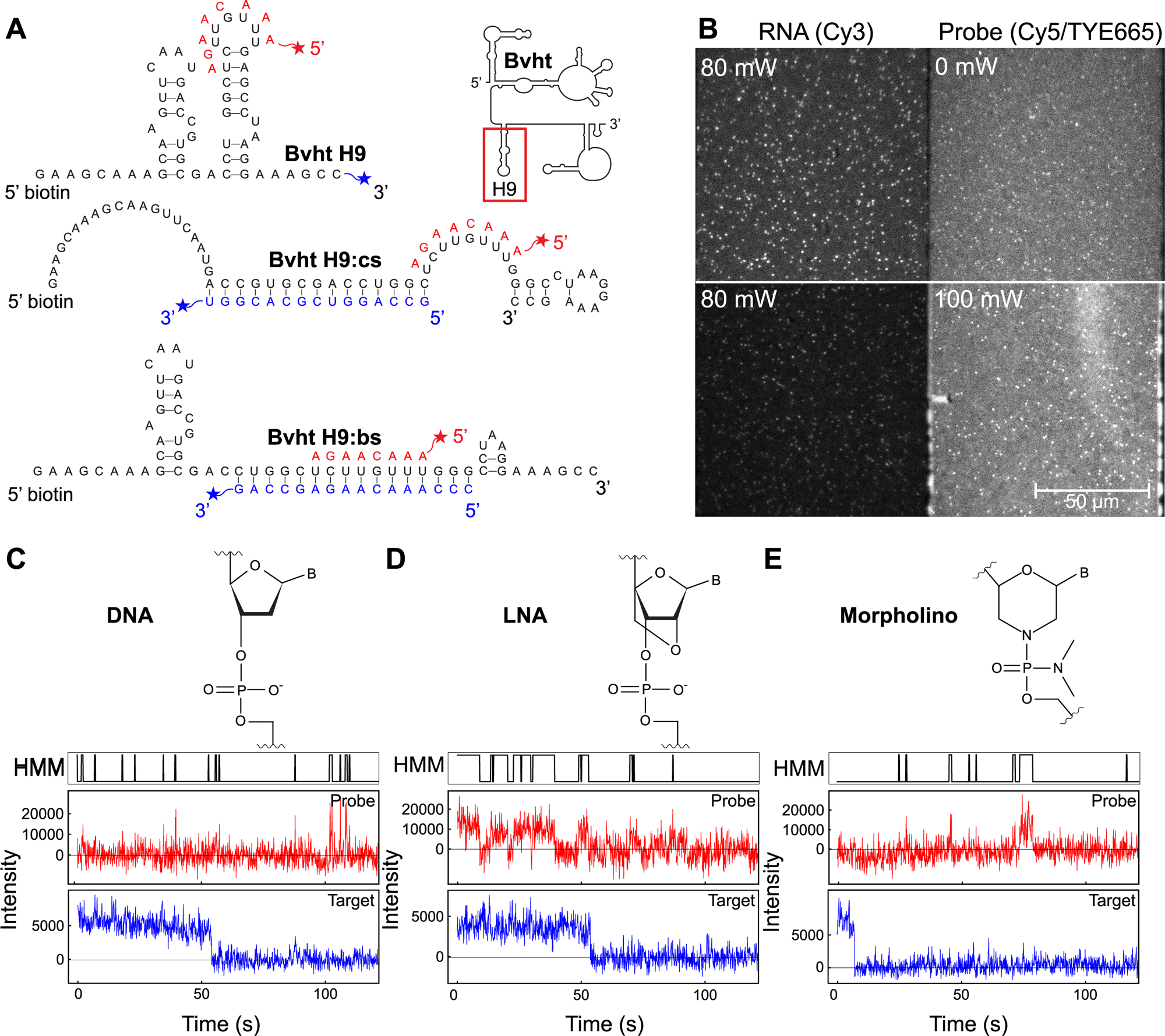
*A*, Model target RNA structures for SiM-KARTS with probe sequence in red adjacent to its binding site. Cy3-labeled complementary RNA strands (cs and bs) stably bound to the primary RNA (Bvht H9) are indicated in blue. Full structure of Bvht and location of H9 region are displayed in the upper right. *B*, Example image from SiM-KARTS data collection. RNA is detected in the left (green) channel via Cy3 fluorescence from either the RNA or a complementary strand (here, 3’ Cy3 cs), while the probe is detected in the right (red) channel via Cy5 or TYE665 fluorescence (here, 5’ TYE665 LNA2). Top: Portion of a field of view under only green laser illumination (source laser power of 80 mW), demonstrating fluorescence from immobilized RNA in the green channel. Spots are visible in the red channel due to spectral bleed-through from Cy3, and diffuse background in the red channel arises from a small amount of direct excitation of freely diffusing probe molecules by the green laser. Bottom: Remaining portion of the same field of view under simultaneous red (source laser power 100 mW) and green (source laser power 80 mW) illumination. Higher diffuse background in the red channel is due to more efficient excitation of freely diffusing probe molecules by the red laser. *C*, Backbone chemistry and example single-molecule trace for a DNA probe binding to Bvht H9:cs. Top: HMM idealization of red (probe) channel trace. Middle: Red (probe) channel trace; Bottom: Green (RNA) channel trace. *D*, Same as *C* for a DNA probe containing one LNA residue binding to Bvht H9:cs. *E*, Same as *C* for a morpholino probe binding to Bvht H9:cs.

## Results

### Experimental Design

A 59-nucleotide sequence containing helix 9 of Bvht (hereafter called “Bvht H9”) was selected from the full-length RNA as a model that allowed different structures to be obtained by hybridizing complementary RNA oligonucleotides (Fig. 1A; sequences in Table 1). Bvht was chosen as an RNA of interest because, in addition to its biological significance, its structure has been well described by chemical probing and SAXS, providing the basis for design of a model RNA for our studies (28, 29). The native structure of Bvht, as previously determined by chemical probing, contains a flexible, single-stranded region to the 5’ side of H9 (28). Using RNAstructure (30), we found that outside the context of the full-length RNA, Bvht H9 is predicted to fold into a structure containing 2 hairpin loops. An 8-nucleotide segment of one loop was chosen as the probe binding site (Fig. 1A). The sequence of this loop is unique within the entirety of Bvht, and therefore is a feasible probing site for studies using the full lncRNA. The complementary strand “cs” was designed to generate a 9-nucleotide single-stranded region around the probing site upon hybridization to Bvht H9, and the blocking strand “bs” was designed to sequester the probing region in double-stranded form (Fig. 1A). Molecules were detected using wide-field prism-based total internal reflection fluorescence (TIRF) microscopy, with Bvht H9 detected in the green emission channel via a Cy3 label on its 3’ end or the 3’ end of cs or bs (ensuring that only hybridized molecules were detected in those cases), and 5’ Cy5- or TYE665-labeled probe detected in the red channel (Fig. 1B). Probes were complementary to the 8-nucleotide target sequence and included one with a fully DNA backbone (“DNA”), one with an LNA residue at position 2 (“LNA2”) in an otherwise DNA backbone, one with LNA residues at positions 2 and 3 in an otherwise DNA backbone (“LNA23”) and one with a morpholino backbone (“MO”) (Fig. 1C-E). The 8-nucleotide length and the number and location(s) of LNA residues were selected to achieve melting temperatures of the probe:RNA hybrid around or below room temperature. This is desirable for SiM-KARTS, as transient, reversible probe binding provides a kinetic signature of the structure of the target RNA without stably occluding the binding site and thereby impacting the structure of the target. Fluorescence intensity traces from the red (probe) channel were idealized using hidden Markov modeling (HMM) to define probe-bound and probe-unbound states (Fig. 1C-E). Ideal probe concentration was determined to be 60 nM by analyzing SiM-KARTS data collected with 20-100 nM LNA2 and comparing changes in bound and unbound dwell times (Fig. S1).

**Table 1.**
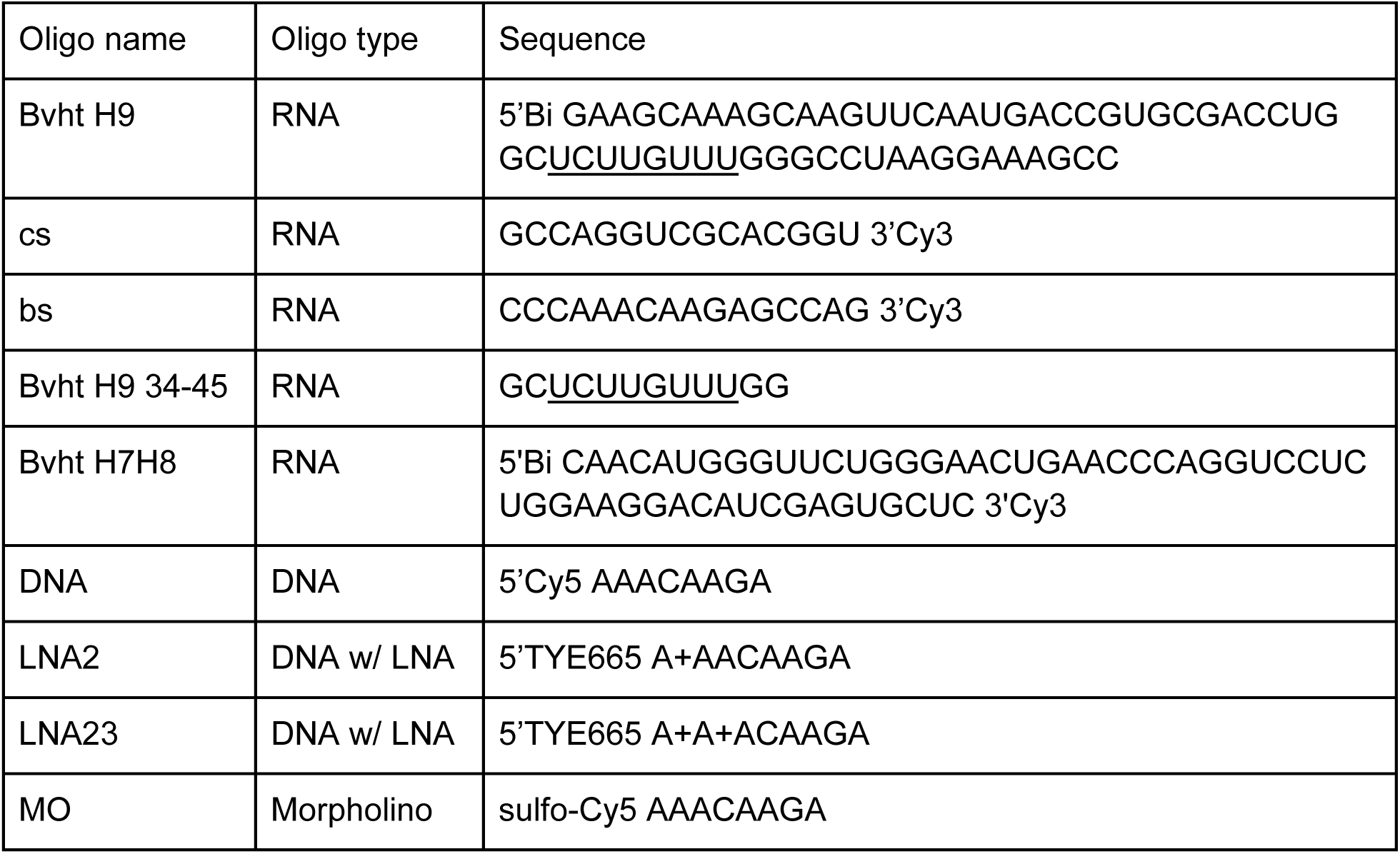
Sequences of oligonucleotides. Probe binding sites are underlined in Bvht H9 and Bvht H9 34-45 sequences. “+X” indicates that nucleotide X is an LNA residue in an otherwise DNA oligonucleotide.

### SiM-KARTS probe chemistry impacts RNA binding properties

We first investigated the impact of probe chemistry on binding behavior to Bvht H9 (containing a partially occluded probe binding site) and Bvht H9:cs (containing a single-stranded probe binding site). The ideal probe will bind in a manner sensitive to structure, and have bound dwell times that are long enough to be readily distinguished from background noise, yet not so long that the binding impacts the structure of the target RNA. The rates of probe binding and dissociation were first quantified using cumulative distribution curves (Fig. 2; additional details in Fig. S2), which show the probability that a bound or unbound segment of a trace has a duration less than or equal to the corresponding time indicated on the x-axis. A slower rise from zero to 1 for unbound dwells indicates that longer intervals between binding events are typical. Similarly, for bound dwells, a slower rise from zero to 1 indicates that longer binding events are typical.

**Figure 2.**
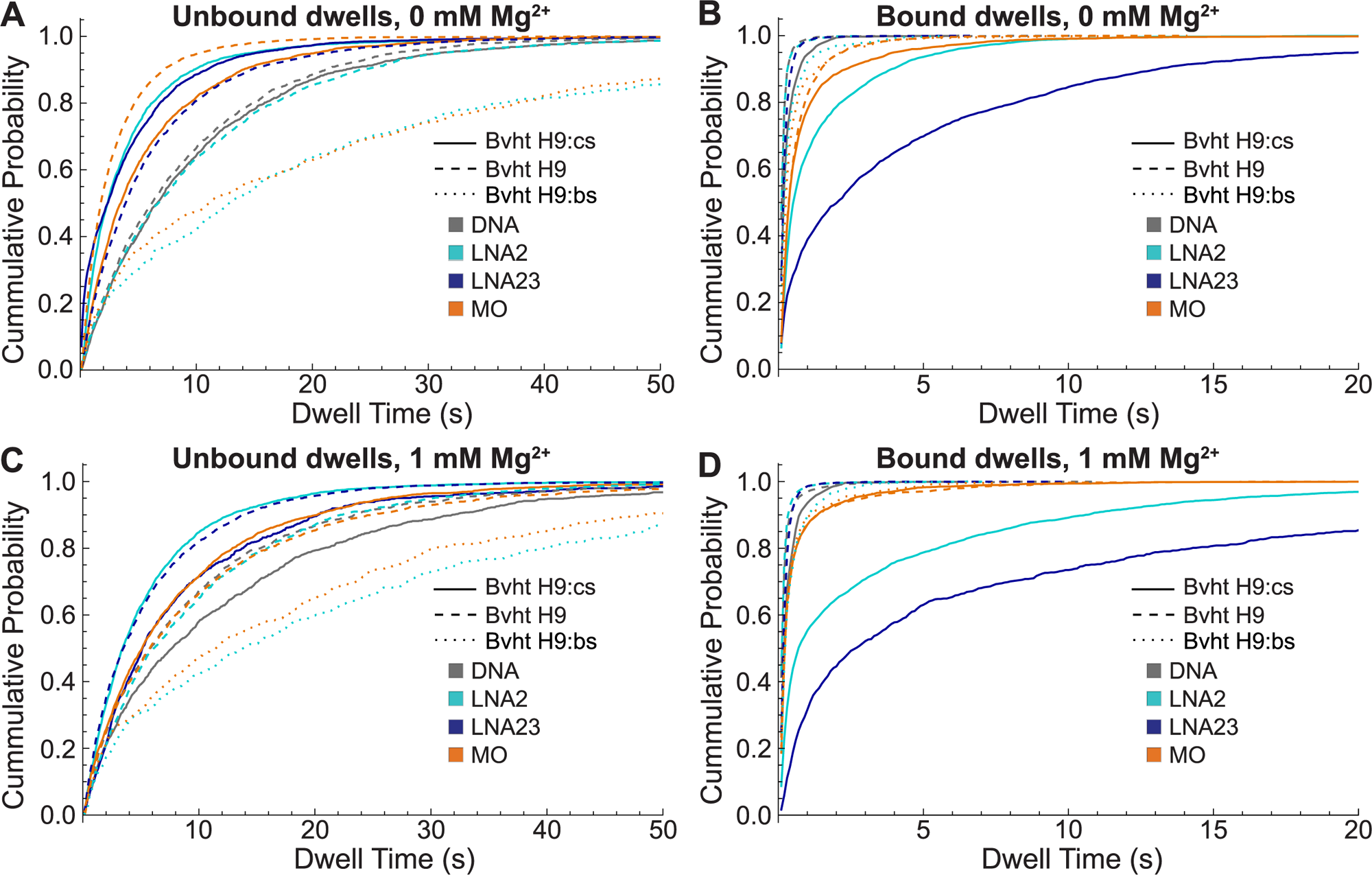
Impact of probe chemistry and Mg^2+^ on bound and unbound dwell times determined by HMM. *A*, Cumulative distributions of unbound dwell times obtained at 0 mM MgCl_2_ for different probe types binding to Bvht H9 (dashed), Bvht H9:cs (solid) or Bvht H9:bs (dotted). Here and throughout, data with DNA probe are plotted in gray, LNA2 in variations of green and cyan, LNA23 in dark blue and MO in variations of orange and brown. *B*, Bound dwell times obtained at 0 mM MgCl_2_ for different probe types binding to Bvht H9, Bvht H9:cs or Bvht H9:bs. *C*, Same as *A* but at 1 mM MgCl_2_. *D*, Same as *B* but at 1 mM MgCl_2_.

Characteristic bound and unbound dwell times were quantified by fitting these curves with single- or double-exponential functions (Fig. S2G-J). Unbound dwell time distributions were well fit (R^2^>0.97) with single-exponential functions in all cases examined, while bound dwell time distributions required double-exponential fits in many cases (Fig. S2I; Tables S1, S2), consistent with previous studies (11). To facilitate direct comparison, bound dwells for all conditions were therefore modeled with double-exponential functions.

The *unbound* dwell times differ for each combination of target RNA structure (Bvht H9 or Bvht H9:cs) and probe chemistry (DNA, LNA2, LNA23 or MO) (Fig. 2A,C; Table 2). Unbound dwell times are longer in the presence of 1 mM Mg^2+^ for all probe types binding to Bvht H9:cs, while Mg^2+^ has little impact on binding to Bvht H9. The DNA probe generally has the longest unbound dwell times across all conditions, though the unbound dwell times for DNA and LNA2 probing Bvht H9 are very similar. LNA2 consistently binds Bvht H9:cs more quickly than Bvht H9, showing strong discrimination between the two structures in both the absence and presence of 1 mM Mg^2+^. LNA23 likewise exhibits faster binding to Bvht H9:cs than Bvht H9 in 0 mM Mg^2+^, but has opposite behavior at 1 mM Mg^2+^.

**Table 2.** Characteristic dwell times of probes binding to Bvht H9 or Bvht H9:cs. τ_unbound_ is obtained from a single-exponential fit to the cumulative distribution of unbound dwell times in Fig. 2. τ_bound_ is presented as a weighted average determined from a double-exponential fit to the cumulative distribution of bound dwell times. Individual component parameters and R^2^ values are reported in Tables S1-S2. The standard deviation σ, obtained from independently fitting three randomly selected subsets of the traces (bootstrapped sets), is noted in parentheses. “N” indicates the number of single-molecule traces included in each condition’s analysis, “D/N” indicates the average number of bound dwells observed per trace, and the total number of bound dwells analyzed is the product of the two.

|  |  | Bvht H9 |  |  |  | Bvht H9:cs |  |  |  |
| --- | --- | --- | --- | --- | --- | --- | --- | --- | --- |
| Probe | [Mg<sup>2+</sup>]<br>(mM) | T<sub>unbound</sub> (s)<br>(σ) | T<sub>bound,avg</sub> (s)<br>(σ) | N | D/N | T<sub>unbound</sub><br>(s) (σ) | T<sub>bound,avg</sub> (s)<br>(σ) | N | D/N |
| DNA | 0 | 9.0 (0.1) | 0.146 (0.002) | 118 | 18 | 9.5 (0.4) | 0.24 (0.04) | 93 | 16 |
| DNA | 1 | 8.9 (0.5) | 0.170 (0.004) | 97 | 16 | 12 (1) | 0.28 (0.02) | 101 | 12 |
| LNA2 | 0 | 9.9 (0.2) | 0.14 (0.01) | 101 | 16 | 3.7 (0.4) | 0.63 (0.09) | 105 | 29 |
| LNA2 | 1 | 9.4 (0.7) | 0.15 (0.01) | 109 | 17 | 5.2 (0.1) | 0.70 (0.05) | 166 | 18 |
| LNA23 | 0 | 5.8 (0.2) | 0.213 (0.004) | 123 | 26 | 3.9 (0.1) | 1.3 (0.4) | 157 | 17 |
| LNA23 | 1 | 5.4 (0.3) | 0.222 (0.004) | 102 | 28 | 8.0 (0.4) | 2.4 (0.5) | 108 | 8 |
| MO | 0 | 7.6 (0.6) | 0.33 (0.02) | 115 | 20 | 5.4 (0.5) | 0.51 (0.03) | 112 | 18 |
| MO | 1 | 9.0 (0.3) | 0.30 (0.02) | 117 | 15 | 7.6 (0.3) | 0.31 (0.01) | 126 | 20 |

It is apparent that the increasing inclusion of LNA residues leads to longer *bound* dwell times, particularly with the target Bvht H9:cs (Fig. 2B,D; Table 2). There is also distinct separation between the bound dwell times of LNA probes for different RNA structures, with longer bound times observed for Bvht H9:cs, which has a more accessible binding site, than Bvht H9.

Comparatively, the bound dwell times for the DNA and MO probes do not discriminate between structures as effectively. Interestingly, the cumulative distributions of bound dwells for Bvht H9:cs are best described with a double-exponential fit, while bound dwells for Bvht H9 can be adequately described using a single exponential for all probes except MO. In these cases, the dominant component of a double-exponential fit (as reported for all bound dwells herein) has a weight of nearly 100% (Table S1). This distinction between target RNA structures is most evident with the LNA probes, where the heavier component is weighted at 99-100% for Bvht H9, and 51-62% for Bvht H9:cs (Table 2). The DNA probe exhibits the same trend but has a less drastic difference between structures, where the dominant component is weighted at 100% for Bvht H9 and the faster component is weighted at 81-88% for Bvht H9:cs.

SiM-KARTS is a high-background measurement due to the presence of freely diffusing fluorescent probe in the sample chamber (Fig. 1B). As a result, we performed extensive negative controls to ensure that the events identified by HMM were dominated by true probe binding rather than noise. Experiments with Bvht H9 hybridized to a blocking strand that completely occludes the binding site (Bvht H9:bs) showed that LNA2 and MO display significantly fewer binding events per trace (∼6) and longer unbound dwell times (∼20 s) than they do with Bvht H9 or Bvht H9:cs as the target (Fig. 2; Table 3), confirming that probe binding is strongly inhibited by competition for occupation of its complementary sequence. Further studies with LNA2 showed that in the absence of any target RNA, or in the presence of a target RNA lacking a complementary binding site (Bvht H7H8), the incidence of apparent probe binding events was lower still at only 1 or 2 bound dwells per trace, respectively, identified by HMM (example traces and quantification in Fig. S3). Similar numbers were observed for Bvht H9:cs with no probe in the sample chamber. Thus, a significant majority of bound dwells identified by HMM in cases where a probe and an accessible complementary sequence are both available (Bvht H9, Bvht H9:cs) represent true probe binding events, as do many of the observed binding events to the occluded Bvht H9:bs binding site. Spurious fluctuations identified as binding events by HMM would have the effect of slightly decreasing the apparent bound and unbound dwell times relative to true probe binding behavior.

**Table 3.** Characteristic dwell times for LNA2 and MO binding to Bvht H9:bs. τ_unbound_ is obtained from a single-exponential fit to the cumulative distribution of unbound dwell times in Fig. 2. τ_bound_ is presented as a weighted average determined from a double-exponential fit to the cumulative distribution of bound dwell times. Individual component parameters and R^2^ values are reported in Table S3. The standard deviation σ, obtained from independently fitting three bootstrapped sets, is noted in parentheses. “N” indicates the number of single-molecule traces included in each condition’s analysis and “D/N” indicates the average number of bound dwells observed per trace.

|  |  | Bvht H9:bs |  |  |  |
| --- | --- | --- | --- | --- | --- |
| Probe | [Mg<sup>2+</sup>] (mM) | T<sub>unbound</sub> (s) ($\sigma$) | T<sub>bound,avg</sub> (s) ($\sigma$) | N | D/N |
| LNA2 | 0 | 20 (2) | 0.29 (0.02) | 103 | 6 |
| LNA2 | 1 | 22 (3) | 0.33 (0.01) | 102 | 5 |
| MO | 0 | 19 (2) | 0.34 (0.01) | 103 | 6 |
| MO | 1 | 18 (2) | 0.29 (0.02) | 100 | 7 |

We also considered whether Förster resonance energy transfer (FRET) could complicate the ability of HMM to identify binding events, given that the Cy3 label used to localize target RNA molecules is separated from the Cy5 or TYE665 label on a bound probe by less than 10 nm. We found that obvious evidence of FRET could be observed by visual inspection only in a small fraction of traces (∼1%) as a decrease in Cy3 fluorescence during periods when the probe was bound. However, Cy3 was active (not photobleached) for only 10% of frames across Bvht H9 traces and 17% of frames across Bvht H9:cs traces, and there was no significant difference in the number of frames when a probe was determined to be bound while Cy3 was active (2.6% of frames and 39% of frames for LNA2 binding to Bvht H9 and Bvht H9:cs, respectively) compared to after photobleaching (2% and 39% of frames for Bvht H9 and Bvht H9:cs, respectively). This indicates that FRET does not impact our analysis of bound and unbound dwell times. Potential impacts of FRET would be mitigated with longer target RNAs where the fluorophore used for localization can be placed farther from the probe binding site.

The precision of our SiM-KARTS measurements was further quantified by comparing dwell times extracted from movies collected on the same day, movies collected across different days (Fig. S4A-B; Table S4), and batches of traces (termed bootstrapped sets) selected randomly from within the combined data across all days (Fig. S4C-D; Table S5) (data on LNA2 binding to Bvht H9:cs). A small amount of variation in unbound and bound dwells is observed in data collected on the same day (Fig. S4A-B), with both quantities differing by 6-7% (standard deviation divided by mean) across movies on the day with higher variability (Table S4).

Comparable variation of ∼5% in unbound dwells was observed across bootstrapped sets (Table S5). In contrast, systematic variation of unbound dwell times is seen in the case of data collected on separate days, with a difference of 32% (difference divided by mean) between data pooled from movies on each day. This is likely due to inconsistencies in the concentration of binding-competent probe across freeze-thaw cycles, which could result from factors such as degradation (though this is unlikely for oligonucleotides with the backbones investigated here), pipetting uncertainty, adherence of the probe to walls of the tube, and gradual precipitation. In the results reported herein, all data for a given probe was collected on the same day to ensure accurate comparisons between target RNA structures and ionic conditions, except for MO, for which differences in unbound dwells should not be necessarily considered significant.

### Melting temperatures confirm increased binding stability of LNA and MO probes

As an orthogonal metric of binding stability, thermal denaturation experiments were performed for each probe type bound to Bvht H9 34-45, a short RNA containing the probing sequence with 2-nucleotide overhangs on either end (Fig. 3). Melting curves were recorded in the absence or presence of 1 mM Mg^2+^ by monitoring absorbance at 260 nm, and were fitted (Fig. S5) to extract melting temperatures (T_M_) and the width of the melting transition (ΔT). As expected, the DNA probe has the lowest T_M_, and the T_M_ values of LNA probes increase with the number of LNA residues incorporated (Table 4). The morpholino probe is similar in melting behavior to LNA2, but shows lower sensitivity to Mg^2+^. The T_M_ obtained for the DNA probe represents an upper limit, as the slope of the melting curve indicates that the probe does not fully hybridize to the RNA even at 0 °C. The T_M_ values of probe hybrids with Bvht H9 34-45 follow the same trends as the bound dwells of probes binding to Bvht H9:cs observed by SiM-KARTS, with higher T_M_ (Table 4) values associated with longer bound dwell times (Table 2). Interestingly, the measured T_M_ values for LNA probes are significantly higher than their estimated values from IDT OligoAnalyzer (31–34). However, the model used by that calculator is experimentally parameterized only for LNA:DNA hybrids, and approximates LNA nucleotides hybridizing with RNA as RNA nucleotides (35). Notably, our results more closely approximate the predictions for LNA:DNA hybrids (Table 4). Additionally, no such melting temperature calculator exists for morpholinos, with most studies of morpholino-nucleic acid hybrids being limited to interactions with DNA only (36, 37). However, the MO:RNA hybrids that have been studied were shown to have higher T_M_ values than corresponding DNA:RNA hybrids by nearly 10 °C (38), which is consistent with the MO:RNA melting temperatures measured here.

**Figure 3.**
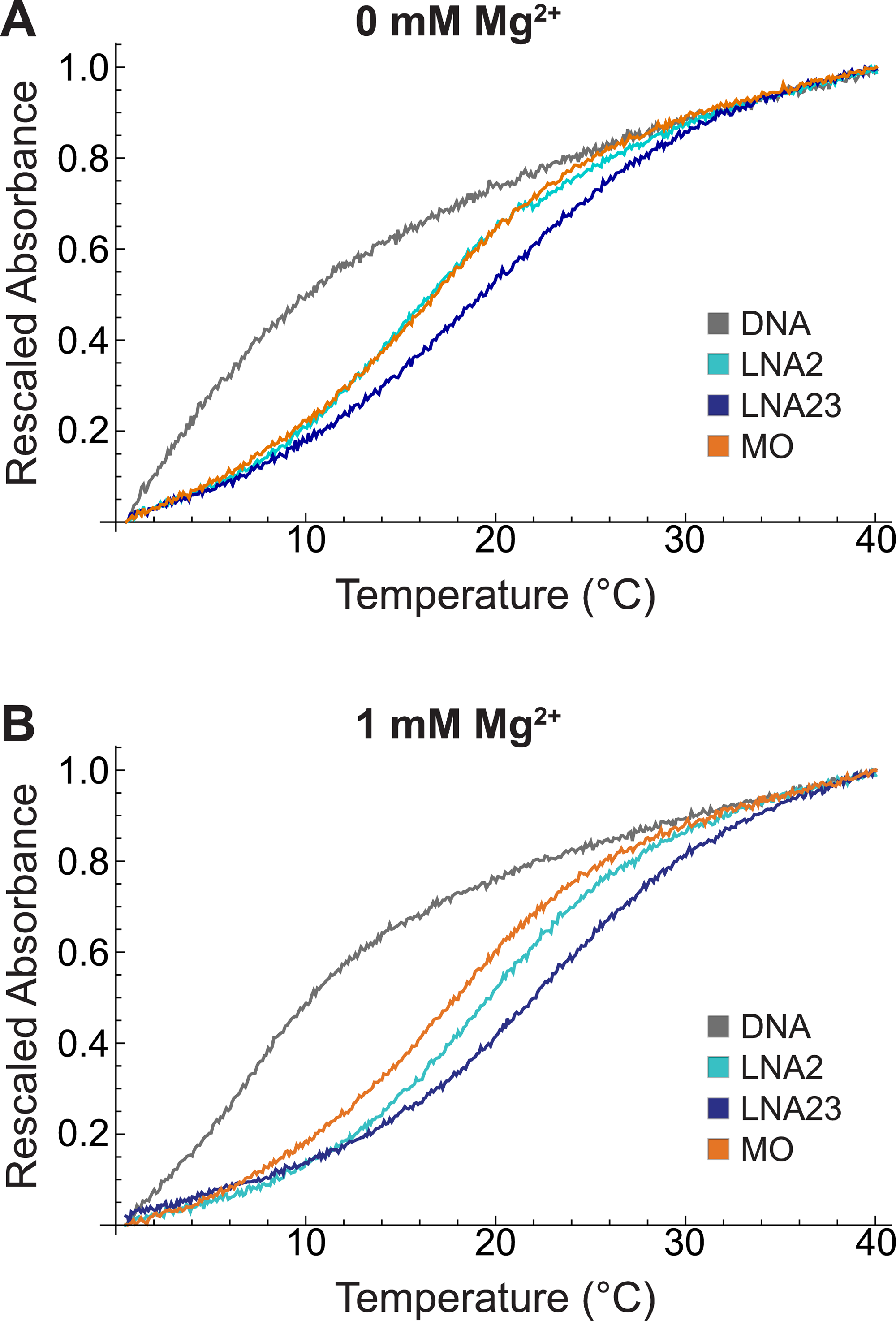
Melting curves for hybrids of each probe type and Bvht H9 34-45 measured by absorbance at 260 nm. For ease of comparison, curves are shifted to begin at an absorbance value of 0 and normalized to end at a value of 1. *A*, Melting curves for each probe type recorded at 0 mM MgCl_2_. *B*, Melting curves for each probe type recorded at 1 mM MgCl_2_.

**Table 4.** Melting temperature (T_M_) and width of the melting transition (ΔT) values for each probe type hybridized to Bvht H9 34-45 in the absence or presence of 1 mM MgCl_2_. Predicted T_M_ values were obtained using IDT OligoAnalyzer. ^1^Upper limit of T_M_ due to incomplete hybridization at 0 °C. ^2^RNA target calculation setting/DNA target calculation setting.

| Probe | [Mg<sup>2+</sup>] (mM) | T<sub>M</sub> from fit (°C) | ΔT from fit (°C) | T<sub>M</sub> predicted (°C) |
| --- | --- | --- | --- | --- |
| DNA | 0 | 6.1<sup>1</sup> | 2.5 | 4.1 |
| DNA | 1 | 7.2<sup>1</sup> | 3.0 | 6.5 |
| LNA2 | 0 | 16.9 | 3.5 | 5.4/18.8<sup>2</sup> |
| LNA2 | 1 | 19.4 | 3.5 | 7.8/21.5<sup>2</sup> |
| LNA23 | 0 | 20.2 | 4.1 | 9/22.4<sup>2</sup> |
| LNA23 | 1 | 20.9 | 3.9 | 11.5/21.5<sup>2</sup> |
| MO | 0 | 17.6 | 3.4 | - |
| MO | 1 | 18.1 | 3.3 | - |

### LNA probes distinguish structures on a trace-by-trace basis

The analyses of cumulative distribution functions described above demonstrated that all probe types exhibit distinguishable dwell time distributions when binding to pure BvhtH9 or pure BvhtH9:cs. However, it is particularly desirable for a probe to enable sorting of individual traces into structural categories within a heterogeneous mixture. To assess the ability of different probe types to distinguish traces arising from one target RNA structure from another, we developed a clustering approach in which each trace was assigned a kinetic signature characterized by the average unbound dwell time ⟨t_unbound_⟩ and the average bound dwell time ⟨t_bound_⟩ observed within it. We visualized the results using scatter plots with each trace represented by a point at its (⟨t_bound_⟩,⟨t_unbound_⟩) coordinates (Fig. 4). We then compared the (⟨t_bound_⟩,⟨t_unbound_⟩) distributions observed with Bvht H9 or Bvht H9:cs as the target RNA for each probe type in the absence and presence of 1 mM MgCl_2_. All probe types yielded distributions for Bvht H9 and Bvht H9:cs whose medians were distinguishable to a high degree of statistical significance (Table 5), with p-values ranging from 7*10^-6^ for MO in 1 mM Mg^2+^ to 2*10^-42^ for LNA23 in 0 mM Mg^2+^ (multivariate Mann-Whitney test). However, the distributions for Bvht H9 and Bvht H9:cs overlap significantly in the cases of DNA and MO (Fig. 4A-B and G-H), suggesting that these probes would not enable traces obtained from a heterogeneous mixture to be assigned to specific structures.

**Figure 4.**
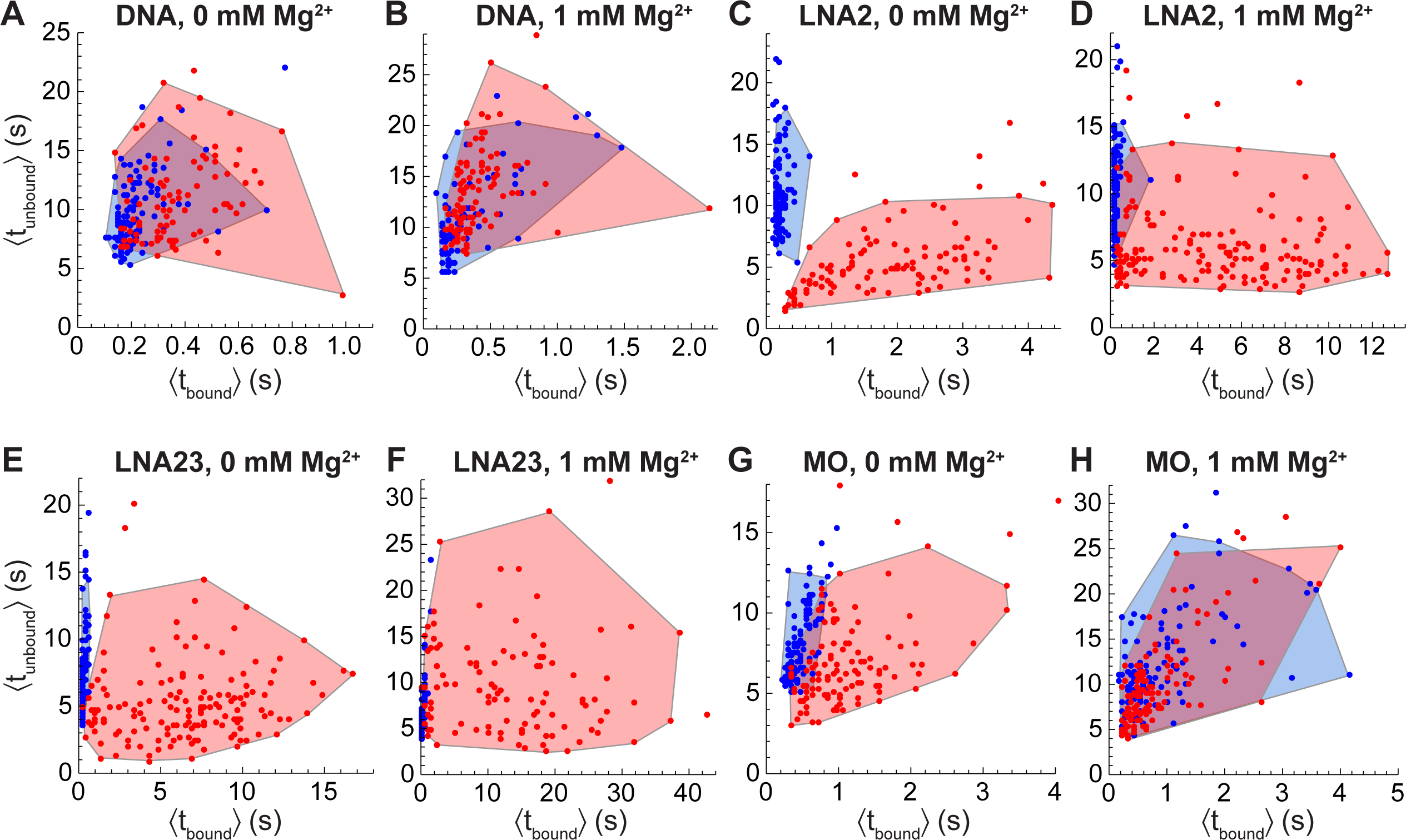
Clustering of traces according to their average bound and unbound dwell times. Each trace is represented by a point located at x-coordinate equal to the average duration of probe-bound segments within it, ⟨t_bound_⟩, and y-coordinate equal to the average duration of unbound segments within it, ⟨t_unbound_⟩. Shaded regions are convex hulls surrounding the 95% of points closest to the centroid, which is located at the median values of ⟨t_bound_⟩ and ⟨t_unbound_⟩. *A*, Scatter plot of ⟨t_unbound_⟩ vs. ⟨t_bound_⟩ and convex hulls for DNA probe binding to Bvht H9:cs (red) or Bvht H9 (blue) at 0 mM MgCl_2_. *B*, Same as *A* but at 1 mM MgCl_2_. *C*, LNA2 at 0 mM MgCl_2_. *D*, LNA2 at 1 mM MgCl_2_. *E*, LNA23 at 0 mM MgCl_2_. *F*, LNA23 at 1 mM MgCl_2_. *G*, MO at 0 mM MgCl_2_. *H*, MO at 1 mM MgCl_2_.

**Table 5.** Quantification of overlap between the (⟨t_bound_⟩,⟨t_unbound_⟩) distributions for Bvht h9 and Bvht H9:cs. ^1^(Median of ⟨t_bound_⟩ across traces, median of ⟨t_unbound_⟩ across traces). ^2^Probability that Bvht H9 and Bvht H9:cs distributions have the same centroid evaluated by the multivariate Mann-Whitney test. ^3^Percentage of Bvht H9 traces whose (⟨t_bound_⟩,⟨t_unbound_⟩) coordinates fall within a convex hull surrounding the Bvht H9:cs (⟨t_bound_⟩,⟨t_unbound_⟩) point cloud. The convex hull omits the 5% of Bvht H9:cs points furthest from the centroid. ^4^Same as ^3^ but with Bvht H9 and Bvht H9:cs reversed.

| Probe | [Mg<sup>2+</sup>]<br>(mM) | Bvht H9<br>centroid (s)<sup>1</sup> | Bvht H9:cs<br>centroid (s)<sup>1</sup> | P-value<sup>2</sup> | % Bvht H9<br>traces within<br>Bvht H9:cs hull<sup>3</sup> | % Bvht H9:cs<br>traces within<br>Bvht H9 hull<sup>4</sup> |
| --- | --- | --- | --- | --- | --- | --- |
| DNA | 0 | (0.2, 9.2) | (0.4,10.4) | 3*10<sup>-15</sup> | 73 | 73 |
| DNA | 1 | (0.2, 9.9) | (0.4,13.5) | 2*10<sup>-8</sup> | 87 | 60 |
| LNA2 | 0 | (0.2, 10.4) | (1.9,4.9) | 1*10<sup>-38</sup> | 0 | 0 |
| LNA2 | 1 | (0.2, 10.3) | (4.5,5.2) | 4*10<sup>-37</sup> | 10 | 6 |
| LNA23 | 0 | (0.3, 6.3) | (6.6,4.8) | 2*10<sup>-42</sup> | 1 | 1 |
| LNA23 | 1 | (0.3, 5.7) | (13.0,8.7) | 1*10<sup>-32</sup> | 2 | 15 |
| MO | 0 | (0.5, 8.0) | (1.0,6.3) | 8*10<sup>-33</sup> | 14 | 43 |
| MO | 1 | (0.6, 10.9) | (0.6,9.0) | 7*10<sup>-6</sup> | 87 | 81 |

We quantified the degree of overlap by constructing a convex hull around the a target RNA’s (⟨t_bound_⟩,⟨t_unbound_⟩) distribution, omitting the 5% of traces with the largest Euclidean distance from the centroid of the distribution (Fig. 4). The degree of overlap was defined as the fraction of one target RNA’s traces whose coordinates fall within the boundary of the other target RNA’s convex hull (by definition, 95% of a target RNA’s traces fall within its own hull). For DNA and MO probes, we observed a high degree of overlap between the (⟨t_bound_⟩,⟨t_unbound_⟩) distributions of Bvht H9 and Bvht H9:cs (Table 5). For example, for DNA in the absence of Mg^2+^, 73% of Bvht H9 traces had (⟨t_bound_⟩,⟨t_unbound_⟩) coordinates within the convex hull of Bvht H9:cs, and 73% of Bvht H9:cs traces fell within the hull of Bvht H9. In contrast, for LNA2 and LNA23 in the absence of Mg^2+^, the percentage of traces falling within the convex hull of the other target RNA was 0% and 1%, respectively. Addition of 1 mM Mg^2+^ increased the overlap between Bvht H9 and Bvht H9:cs distributions for LNA2, LNA23 and MO, though LNA2 and LNA23 retained strong discrimination between the two (for example, only 10% of Bvht H9 traces fell within the convex hull of Bvht H9:cs for LNA2 at 1 mM Mg^2+^). This suggests that probes containing LNA nucleotides could enable categorization of traces obtained from a heterogeneous mixture with a much higher degree of precision than DNA probes.

To test this idea, we divided the 95% of traces closest to the centroid into two pools (termed the “basis” pool and the “test” pool) and generated convex hulls using the traces in the basis pool. We then tested whether the (⟨t_bound_⟩,⟨t_unbound_⟩) coordinates of each trace in the test pool localized within only the correct hull (e.g. Bvht H9 test traces within the Bvht H9 basis trace hull), only the incorrect hull (e.g. Bvht H9 test traces within the Bvht H9:cs basis trace hull), a region where the hulls overlapped, or outside of both hulls entirely. For LNA2 in the absence of Mg^2+^, 85% of the test pool of Bvht H9 traces correctly localized only within the Bvht H9 basis pool’s hull, while none localized within the hull generated from the Bvht H9:cs basis pool (mean across 50 random allocations of traces into basis and test pools). The remaining 15% localized outside of both hulls, but this is expected due to the random allocation of outliers (which increase the footprint of the convex hull) into the basis and test pools, and could be mitigated by accumulating a larger basis pool. Conversely, 85% of Bvht H9:cs test pool traces correctly localized only within the Bvht H9:cs basis pool’s hull and the remaining 15% localized outside of both. LNA23 exhibited almost identical behavior to LNA2 in its ability to discriminate between Bvht H9 and Bvht H9:cs traces. In contrast, for DNA in the absence of Mg^2+^, only 29% of Bvht H9 test pool traces correctly localized only within the Bvht H9 hull, but 57% localized within both hulls due to the extensive overlap between them. Conversely, 27% of Bvht H9:cs traces correctly localized within only the Bvht H9:cs hull while 55% localized within both hulls. These results confirm that LNA probes offer the potential for trace-by-trace sorting into different structural categories, a unique capability among the probe types tested.

### Divalent cations impact probe binding to Bvht H9:cs

The changes in T_M_ observed with and without 1 mM Mg^2+^ show a clear relationship between divalent cation concentration and probe:RNA hybrid stability. To investigate the impact of ionic conditions in SiM-KARTS, we chose to further study the binding of LNA2 and MO, as both showed promise as more stably-binding probes than DNA (Fig. 2; Table 2). For LNA2, the unbound dwell times for the more accessible Bvht H9:cs increase as Mg^2+^ concentration increases, with the less accessible Bvht H9 having consistent unbound dwell times regardless of Mg^2+^ concentration (Fig. 5A, Table 6). Unbound and bound dwell times display little dependence on Na^+^ concentration for either target RNA (Fig. 5C-D). Bound dwells for Bvht H9 are short regardless of Mg^2+^ or Na^+^ concentration (Fig. 5B,D; Table 6). Bound dwells lengthen for Bvht H9:cs between 0 and 1 mM Mg^2+^, but shorten substantially with the addition of 20 mM Mg^2+^, which has previously been used for SiM-KARTS (11), due to a significant increase in the weight of the faster component of the double-exponential fit (Table S7). This suggests that at high concentrations, Mg^2+^ is no longer just stabilizing the interaction between Bvht H9:cs and LNA2, but may be impacting the overall structure of Bvht H9:cs, affecting the accessibility of the probe binding site.

**Figure 5.**
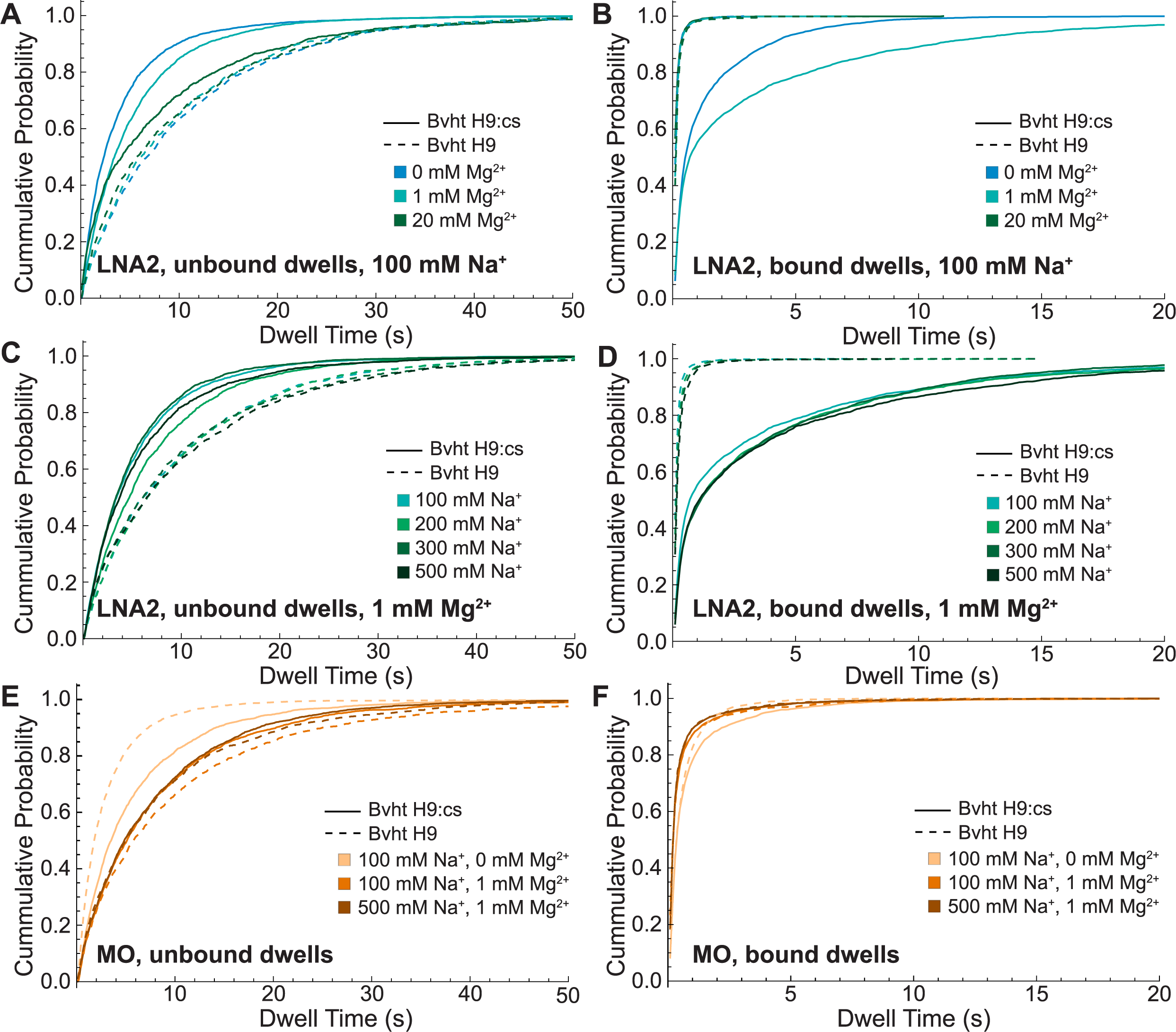
Impact of ionic concentration on dwell times of LNA2 or MO binding to Bvht H9 (dashed) or Bvht H9:cs (solid). *A*, Cumulative distributions of unbound dwell times for LNA2 at 100 mM NaCl and varying MgCl_2_ concentration. *B*, Bound dwell times for LNA2 at 100 mM NaCl and varying MgCl_2_ concentration. *C*, Unbound dwell times for LNA2 at 1 mM MgCl_2_ and varying NaCl concentration. *D*, Bound dwell times for LNA2 at 1 mM MgCl_2_ and varying NaCl concentration. *E*, Unbound dwell times for MO at indicated combinations of NaCl and MgCl_2_ concentration. *F*, Bound dwell times for MO at indicated combinations of NaCl and MgCl_2_ concentration.

**Table 6.** Characteristic dwell times of LNA2 or MO binding to Bvht H9 or Bvht H9:cs in indicated salt concentrations. τ_unbound_ is obtained from a single-exponential fit to the cumulative distribution of unbound dwell times in Fig. 5. τ_bound_ is presented as a weighted average determined from a double-exponential fit to the cumulative distribution of bound dwell times. Individual component parameters and R^2^ values are reported in Table S6-S9. The standard deviation σ, obtained from independently fitting three bootstrapped sets, is noted in parentheses. “N” indicates the number of single-molecule traces included in each condition’s analysis and “D/N” indicates the average number of bound dwells observed per trace. ^1^Data in these rows reproduced from Table 2 for ease of comparison.

|  |  |  | Bvht H9 |  |  |  | Bvht H9:cs |  |  |  |
| --- | --- | --- | --- | --- | --- | --- | --- | --- | --- | --- |
| Probe | [Mg<sup>2+</sup>]<br>(mM) | [Na<sup>+</sup>]<br>(mM) | T<sub>unbound</sub><br>(s) (σ) | T<sub>bound,avg</sub><br>(s) (σ) | N | D/N | T<sub>unbound</sub><br>(s) (σ) | T<sub>bound,avg</sub><br>(s) (σ) | N | D/N |
| LNA2 | 0<sup>1</sup> | 100 | 9.9<br>(0.2) | 0.14<br>(0.01) | 101 | 16 | 3.7<br>(0.4) | 0.63<br>(0.09) | 105 | 29 |
|  | 1<sup>1</sup> | 100 | 9.4<br>(0.7) | 0.15<br>(0.01) | 109 | 17 | 5.2<br>(0.1) | 0.70<br>(0.05) | 166 | 18 |
|  | 1 | 200 | 9.6<br>(0.2) | 0.19<br>(0.01) | 93 | 16 | 6.8<br>(0.5) | 0.9 (0.1) | 157 | 15 |
|  | 1 | 300 | 9.3<br>(0.3) | 0.21<br>(0.01) | 105 | 16 | 5.0<br>(0.2) | 0.89<br>(0.01) | 146 | 18 |
|  | 1 | 500 | 10.0<br>(0.7) | 0.22<br>(0.03) | 103 | 15 | 5.7<br>(0.6) | 1.0 (0.4) | 135 | 15 |
|  | 20 | 100 | 9.1<br>(0.6) | 0.169<br>(0.004) | 89 | 16 | 7.4<br>(0.3) | 0.11<br>(0.02) | 62 | 19 |
| MO | 0<sup>1</sup> | 100 | 7.6<br>(0.6) | 0.33<br>(0.02) | 115 | 20 | 5.4<br>(0.5) | 0.51<br>(0.03) | 112 | 24 |
|  | 1<sup>1</sup> | 100 | 9.0<br>(0.3) | 0.30<br>(0.02) | 117 | 15 | 7.6<br>(0.3) | 0.31<br>(0.01) | 126 | 18 |
|  | 1 | 500 | 7.7<br>(0.4) | 0.30<br>(0.02) | 113 | 18 | 7.5<br>(0.7) | 0.301<br>(0.004) | 135 | 20 |

To further understand the impact of Mg^2+^ concentration on the nature of the binding site encountered by a probe, circular dichroism (CD) spectroscopy was performed on Bvht H9 34-45 and Bvht H9:cs at 0, 1, and 20 mM Mg^2+^. Differences in the structure of Bvht H9 34-45 can be seen in the presence and absence of Mg^2+^, but there are no appreciable differences between 1 mM and 20 mM Mg^2+^ (Fig. 6A). The lineshape of the CD spectrum of Bvht H9:cs remains consistent across Mg^2+^ concentrations, indicating that Mg^2+^ does not induce changes in its global architecture (Fig. 6B). However, the magnitude of the CD signal increases continuously with increasing Mg^2+^, suggesting that this assembly adopts a more ordered structure in the presence of Mg^2+^. This ordered structure may poise the target site in a configuration that is not amenable to probe binding, explaining the slower binding and more rapid dissociation observed at 20 mM Mg^2+^.

**Figure 6.**
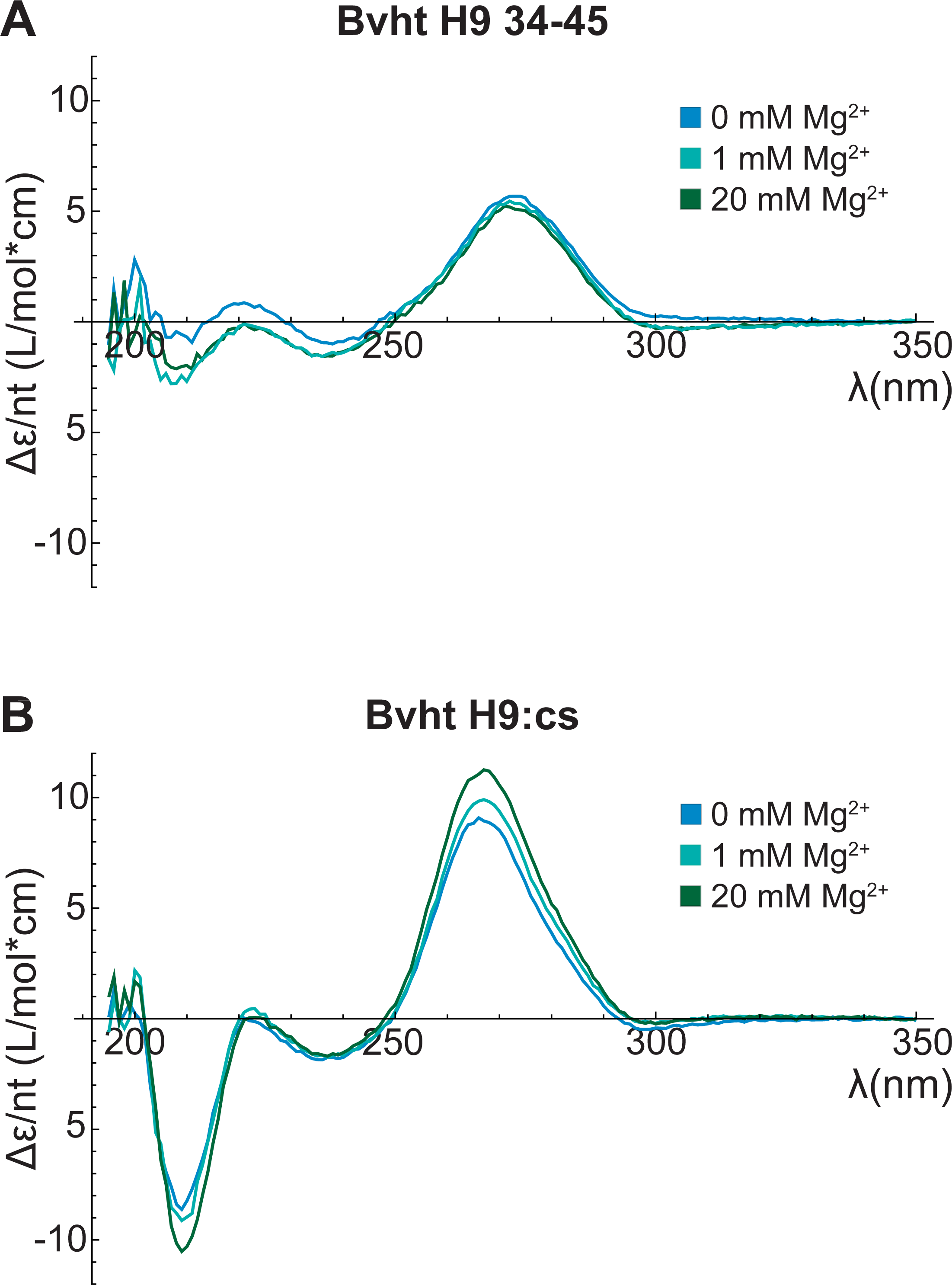
CD spectroscopy of RNA at varying MgCl_2_ concentration. *A*, CD (expressed as Δε per nucleotide) of Bvht H9 34-45 at indicated MgCl_2_ concentrations. *B*, CD of Bvht H9:cs at indicated MgCl_2_ concentrations.

The binding of the morpholino probe was also of particular interest, as its neutral backbone could limit the effect that ions have on probe binding. Here, some variation is seen in the length of unbound dwells, without a clear trend between dwell time and structure probed or salt concentration (Fig. 5E-F; Table 6). The bound dwells are of consistent length and the fit components are of consistent weight for MO binding to Bvht H9 and Bvht H9:cs under all ionic conditions tested. This suggests that the binding stability of the morpholino probe is indeed less sensitive to ionic conditions, but, as noted earlier, is also less sensitive to the structure of the target RNA.

## Discussion

RNA studies using SiM-KARTS have been limited to date, in part due to the optimization necessary to yield clear, interpretable signals from probe binding events. In this work, we found that probes with backbone chemistries beyond DNA can successfully report on RNA structure via SiM-KARTS. Specifically, we found that the incorporation of LNA residues in a DNA oligonucleotide leads to clearer discrimination between target RNA structures in which the probe binding sites exhibit different levels of accessibility. For LNA probes, bound dwell times were notably longer when probing a region that is fully single-stranded and accessible (as in Bvht H9:cs), than they were when probing a partially occluded region (as in Bvht H9). Although the bound dwell times of LNA probes on partially occluded regions were comparable to those observed for LNA2 on the fully occluded Bvht H9:bs complex, the latter was distinguished by much longer unbound dwell times. SiM-KARTS measurements utilizing LNA probes can therefore effectively discriminate among fully accessible, partially occluded and fully occluded binding sites on our model RNA. The optimal number and location of LNA residues is likely to depend on probe length and sequence. We show that with currently available models, melting temperatures predicted with a DNA target setting provide a starting point that can be used to guide design of LNA probes.

Previously, much of the interpretation of SiM-KARTS data was based on *unbound* dwell times being longer for structures with less accessible probing regions, compared to more accessible structures (11–13). Interestingly, we found that this is not necessarily the case under all circumstances. In the absence of Mg^2+^, the dependence of unbound dwell times on the structure of the target RNA was inconsistent across different probe types. Here, the DNA and MO probes demonstrated longer unbound dwells for the more accessible target RNA structure than the less accessible structure, which is contrary to typical interpretation of data. However, in the presence of 1 mM Mg^2+^, LNA and MO probes exhibited longer unbound dwells with the less accessible target structure than the more accessible structure, as may be expected. The unbound dwell times of the DNA probe did not adequately differentiate the two structures.

We have found that *bound* dwell times can also be used to discern information about the structure of the target RNA. There is a clear target structure-dependence of the bound dwell times when using LNA probes, with longer bound dwell times for the structure with a more accessible probe binding site in both the absence or presence of 1 mM Mg^2+^. In the case of DNA, bound dwell times are 50-100% higher for the more accessible structure, but both are very short and are therefore difficult to distinguish from background noise. The morpholino probe, however, exhibits no appreciable difference between structures. We also found that additional information about target RNA structure can be gathered from the component weights of double-exponential fits to bound dwell time distributions. A single component is vastly dominant for DNA and LNA probes binding to more structured target RNA sites, such as Bvht H9 under all conditions and, to a lesser extent, Bvht H9:cs in 20 mM Mg^2+^. In contrast, both components show significant weight for less structured target RNA such as Bvht H9:cs. While the exact mechanism underlying these differences is not known, it is clear that different target structures exhibit different fitting characteristics.

In addition to our analysis of the overall distribution of unbound and bound dwell times observed across many traces, we have found that it is possible, specifically when using LNA probes, to categorize the structures that give rise to individual traces. With DNA and MO probes, significant overlap exists in the behavior observed within individual traces with Bvht H9 and Bvht H9:cs as targets. LNA probes were found to show a higher degree of separation between distributions, allowing individual traces to be assigned to structural categories with higher accuracy. This accuracy is achieved by constructing a kinetic fingerprint from a trace’s bound *and* unbound dwell times, further highlighting the value of considering both quantities holistically.

Previous studies with SiM-KARTS have utilized Mg^2+^ concentrations above those generally considered physiological (39), ranging from 5 to 20 mM (11, 12, 20), to aid in stabilization of probe binding. We have found that for LNA probes, adjusting divalent cation concentration impacts both bound and unbound dwell times; however the impact on bound dwell times is not monotonic as Mg^2+^ concentration increases. Addition of Mg^2+^ increases the stability of the probe:RNA hybrid, as demonstrated by measured melting temperatures, but can also lead to structural changes in the target RNA, as observed by CD. While the exact nature of these conformational rearrangements is not known for Bvht H9, they likely play a role in the non-monotonic dependence of bound dwell times on Mg^2+^ concentration. Changes in monovalent salt concentration do not lead to significant changes in bound or unbound dwells.

Based on these results, we propose that to best optimize probe binding in SiM-KARTS experiments, it is advantageous to change the probe’s chemistry rather than tune binding using environmental variables, such as salt conditions, or by increasing the probe’s length, which may lead to reduced precision for targeting specific regions of interest. Additionally, it is worthwhile to consider both bound and unbound dwell times as well as their exponential fitting characteristics, to gain maximal information about the structure of the binding site and discriminate between similar structures. The probe design and data analysis approaches demonstrated here using a small, controllable model system will facilitate the application of SiM-KARTS to larger RNA targets of unknown structure.

### Experimental procedures Oligonucleotides

To aid with oligonucleotide design, secondary structures of Bvht H9 alone and with potential complementary strand bound were predicted using RNAstructure (30). Bvht H9, Bvht H7H8, and the complementary strands cs and bs (Table 1) were purchased from Dharmacon, with 3’ Cy3 labeling for cs and bs. Bvht H9 was purchased with 5’ biotin, and was 3’ end-labeled with Cy3 hydrazide (Lumiprobe) following established protocols (17) for experiments performed on this RNA without any complementary strand. Bvht H7H8, which has no more than three consecutive bases complementary to the probes used in this study throughout its sequence, was purchased with 5’ biotin and 3’ Cy3 labeling. Probes were designed to be complementary to nucleotides 36-43 of Bvht H9, and the 8-nucleotide length was chosen to achieve a low melting temperature, ensuring transient binding. The DNA probe was purchased from IDT with 5’ Cy5 labeling and DNA/LNA chimera probes were purchased from Qiagen with 5’ TYE665 labeling. TYE665 is a proprietary Cy5 substitute available from Qiagen, which does not offer Cy5 as an LNA modification. Per manufacturer recommendations for chemically modified oligonucleotides of these lengths, all of the above oligonucleotides were HPLC-purified, desalted and, for RNA, deprotected by the manufacturer. The morpholino probe was purchased from GeneTools with 5’ sulfo-Cy5 labeling and was purified by filtration and precipitation by the manufacturer, per their recommendation for oligonucleotides of this type. For melting curves and CD, a truncated version of Bvht H9 was purchased from IDT, containing only the probing region with 2-nucleotide overhangs (Bvht H9 34-45).

### RNA preparation for SiM-KARTS

For single-molecule experiments, 1.5 μM Bvht H9 or Bvht H7H8 was heated to 90 °C in a buffer consisting of 50 mM Tris-HCl and 100 mM NaCl at pH 7.5 for 2 minutes. The RNA was then slowly cooled in air for 10 minutes before being placed on ice. For Bvht H9:cs or Bvht H9:bs, Bvht H9 was prepared at 1 μM in the presence of equimolar cs or bs then annealed as above. A 10 nM dilution of the RNA was prepared shortly before use in experiments.

### SiM-KARTS data collection

Microscope slides were cleaned and passivated using the established DDS-Tween20 method (40). Briefly, quartz slides were cleaned and coated with dichlorodimethylsilane (Sigma Aldrich), then assembled into flow cells, which were stored at 4 °C for up to 3 weeks. Just prior to imaging, passivation was completed by incubating 0.2 mg/mL biotinylated BSA (ThermoFisher) for 5 minutes, 0.2% v/v Tween-20 (Sigma Aldrich) for 10 minutes, and 0.2 mg/mL streptavidin (ThermoFisher) for 5 minutes, each in a buffer consisting of 10 mM Tris-HCl and 50 mM NaCl at pH 8. RNA was flowed on at a concentration of 10 pM in a buffer consisting of 50 mM Tris-HCl and 100 mM NaCl at pH 7.5 and allowed to incubate for 5-10 minutes until an adequate density of molecules, approximately 0.04 spots per µm^2^, was achieved. Probes were flowed on in a buffer consisting of 50 mM Tris-HCl pH 7.5 and variable salt concentrations as specified in figures (100-500 mM NaCl and 0-20 mM MgCl_2_), supplemented with 5 nM 3,4-dihydroxybenzoic acid (Sigma Aldrich), 5 mM Trolox (Sigma Aldrich), and 50 nM Protocatechuate 3,4-dioxygenase (MP Biomedicals) as an oxygen scavenging system. Probes were present at 60 nM unless otherwise noted. This mixture was incubated in the sample chamber for 10 minutes prior to data acquisition. Data were collected at room temperature on a home-built prism-type total internal reflection fluorescence microscope (Leica DMi8 inverted microscope, 63x oil immersion objective). Fluorophores were excited continuously during data acquisition using a 532 nm laser (Coherent OBIS 532 LS) at 80 mW and a 637 nm laser (Coherent OBIS 637 LX) at 100 mW. Emission was split into red and green channels (Chroma T635lpxr longpass dichroic beamsplitter) and cleanup filters were placed in both channels (Chroma ET655lp in the red channel, Chroma ET585/65m in the green channel, and ZET635NF and ZET532NF in both channels). The channels were projected side-by-side onto an Andor iXon 888 EMCCD camera. Movies were 2,000 frames in duration and were recorded with an exposure time of 100 ms and an EM gain setting of 200.

### SiM-KARTS data analysis

Coordinates of RNA molecules in the green channel were identified for each video file using the FIJI “find maxima” feature after rolling ball background subtraction. The tolerance for peak finding was defined for each movie by eye, taking into account the difference between signal and background. Custom MATLAB code was then used to extract fluorophore intensity traces for these locations and the corresponding locations in the red channel, which were identified by performing a calibration using fluorescent microspheres (FluoSpheres, 0.2 μm, red, carboxylate-modified microspheres, ThermoFisher) visible in both channels. Traces were corrected for spectral bleed-through of Cy3 emission into the red channel and were manually selected for further processing using custom MATLAB code if they exhibited single-step photobleaching of Cy3 and a flat noise floor in the red channel. For experiments with no immobilized RNA, traces were extracted from periodically distributed locations throughout the field of view and ones with a flat baseline in the red channel were selected for further analysis. Data from 2 to 3 movies taken on the same day were combined to yield the number of traces indicated by “N” in each data table. Selected traces were placed through a Wiener filter with a range of 1 to reduce background noise and then normalized to each have a noise floor of zero and a peak value of 2. Filtered traces underwent hidden Markov modeling in QuB Online software (41–45) using a 2-state model (probe bound and unbound) to obtain idealized traces. Starting parameter guesses for HMM were typically amplitudes of 0.2 and 1 with standard deviations of 0.2 and 0.6 for the unbound and bound states, respectively, and rates of k_on_=0.3 s^-1^ and k_off_=1 s^-1^. The algorithm was typically allowed to re-estimate all of these parameters except for the standard deviation of the signal in the unbound state. Starting guesses and parameters allowed to float were altered in cases of obvious under- or over-fitting.

Transition rates were analyzed by obtaining the durations of dwells in the bound and unbound states, omitting segments that were truncated by the beginning or end of the trace. The cumulative distributions of each were fit with single- or double-exponential functions in Mathematica (Wolfram Alpha) (Eq. 1 or Eq. 2, respectively, where w_i_ is the weight of component i and τ_i_ is its time constant).

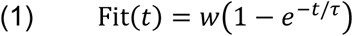

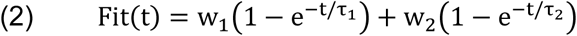

To obtain uncertainties for dwell times, traces were randomly split into three pools (termed “bootstrapped sets”), and the bound and unbound dwell times observed within each set were independently assembled into cumulative distribution curves and fitted. Tables 2, 3, 7 and 8 present the mean and standard deviation of the reported fitting parameters across the three bootstrapped sets.

Scatter plots were generated and analyzed in Mathematica. The durations of all bound segments of a trace were averaged together and the durations of all unbound segments were averaged together, omitting segments truncated by the beginning or end of the trace, to generate that trace’s (⟨t_bound_⟩,⟨t_unbound_⟩) coordinates. Scatter plots were constructed with each trace represented by a point at its (⟨t_bound_⟩,(⟨t_unbound_⟩) coordinates. The coordinates of the centroid were identified as (median of ⟨t_bound_⟩ across traces, median of ⟨t_unbound_⟩ across traces). The medians of Bvht H9 and Bvht H9:cs (⟨t_bound_⟩,⟨t_unbound_⟩) distributions were compared using the multivariate Mann-Whitney test, which is appropriate for data with multiple dependent variables (here, ⟨t_bound_⟩ and ⟨t_unbound_⟩) that are not normally distributed. A convex hull was constructed around the points using the Mathematica built-in function “ConvexHullMesh”, omitting the 5% of points with the largest Euclidian distance from the centroid. Each trace’s coordinates were tested for whether they fell inside a dataset’s convex hull using the Mathematica built-in function “RegionMember”. To quantify the accuracy with which traces could be categorized, the 95% of traces closest to the centroid were randomly assigned to the basis pool or test pool (half in each pool), a convex hull was constructed around the coordinates of the basis pool, and the coordinates for traces in the test pool were queried using “RegionMember” to determine which hulls they localized within. The reported results are the average over 50 independent randomizations.

### Melting curves

Melting curves were recorded on a Jasco J-1500 CD spectrometer in a low headspace, 1 cm pathlength cuvette (Starna 26.100/LHS). 3 μM each of probe and Bvht H9 34-45 were combined in 20 mM sodium phosphate (pH 7.4) supplemented with NaCl to attain a total Na^+^ concentration of 100 mM and MgCl_2_ at 0 or 1 mM, as specified in figures. While Tris buffer was used for SiM-KARTS experiments to match earlier studies (11, 13), phosphate buffer was preferred for melting curves and CD spectroscopy due to the milder temperature-dependence of its pH and its superior ultraviolet transparency. Samples were incubated at 0 °C for 10 minutes before being heated at 0.2 °C/min from 0 °C to 40 °C, with absorbance monitored at 260 nm. A buffer scan was subtracted, and the resulting curves were shifted to coincide with zero at low temperature and normalized to end at 1 for display. T_M_ values were obtained by fitting the curves in Mathematica based on a previously described procedure (46). First, the high-temperature segment of the curve with constant slope was fitted with a line, which was extrapolated to lower temperature. The fraction of strands that were hybridized was calculated using Eq. 3:

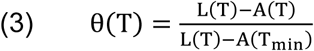

where L(T) is the value of the linear fit at temperature T, A(T) is the absorbance at temperature T, and T_min_ is the minimum temperature measured. θ(T) was then fitted using Eq. 4:

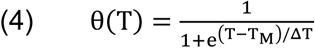

where T_M_ is the melting temperature and ΔT is the width of the melting transition. T_M_ values were predicted using the Integrated DNA Technologies Oligo Analyzer (https://www.idtdna.com/pages/tools/oligoanalyzer) (31) using an oligo concentration setting of 1.5 μM (following recommendations for equimolar strand concentrations) and ion concentrations to match desired conditions.

### Circular dichroism spectroscopy

CD spectra were recorded on a Jasco J-1500 CD spectrometer in a 1 cm pathlength cuvette (Starna 26.100F-Q-10/Z15). Bvht H9:cs or Bvht H9 34-45 was prepared at an optical density of ∼0.8 in a buffer consisting of 20 mM sodium phosphate (pH 7.4) supplemented with NaCl to attain a total Na^+^ concentration of 100 mM and MgCl_2_ at 0 or 1 mM, as specified in figures. Bvht H9:cs was then annealed as described above by heating to 90 °C for 2 minutes and then slowly cooling in air for 10 minutes. Spectra were collected from 350-195 nm at 20 °C, at a 10 nm/min scanning speed. Spectra were processed by subtracting a buffer scan, zeroing the long-wavelength baseline, and converting to Δε per nucleotide using Eq. 5, where CD is the CD signal in millidegrees, L is the pathlength, C is the RNA concentration determined by A260, and n is the number of nucleotides in the RNA.

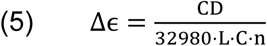

## Data availability

Data and custom software underlying this report are available upon request to the corresponding author

## Supporting information

This article contains supporting information.

## Author contributions

KRP: Conceptualization, formal analysis, investigation, methodology, visualization, writing-original draft. AMM: Formal analysis, investigation, writing - review & editing. MK: Formal analysis, investigation, writing - review & editing. JRW: Conceptualization, formal analysis, funding acquisition, project administration, supervision, visualization, writing - review & editing

## Funding and additional information

This work was supported by American Heart Association grants 857651 (to JRW) and 23IAUST1029376 (to Daniel Grimes), National Institutes of Health grant R35 GM147229 (to JRW), and National Science Foundation Graduate Research Fellowship Program under Grant No. 2236419 (to KRP). The content is solely the responsibility of the authors and does not necessarily represent the official views of the National Institutes of Health. Any opinions, findings, and conclusions or recommendations expressed in this material are those of the author(s) and do not necessarily reflect the views of the National Science Foundation.

## Abbreviations

The abbreviations used are: lncRNA, long non-coding RNA; SiM-KARTS, single-molecule kinetic analysis of RNA transient structure; smFRET, single-molecule Förster resonance energy transfer; SD, Shine Dalgarno; SAXS, small angle x-ray scattering; LNA, locked nucleic acid; MO, morpholino; Bvht, *Braveheart*; FRET, Förster resonance energy transfer; T_M_, melting temperature; CD, circular dichroism.

