## Supporting Information for "Alternative probe chemistries for single-molecule analysis of long non-coding RNA"

**Contents**

Figure S1. Impact of probe concentration on dwell times of LNA2 binding to Bvht H9:cs
Figure S2. Overview of dwell time distribution analysis
Figure S3. Representative single-molecule traces of control conditions
Figure S4. Cumulative distribution plots illustrating SiM-KARTS data reproducibility
Figure S5. Processed melting curves and fits
Table S1. Detailed fitting results for probes binding to Bvht H9
Table S2. Detailed fitting results for probes binding to Bvht H9:cs
Table S3. Detailed fitting results for probes binding to Bvht H9:bs
Table S4. Detailed fitting results for experimental replicates of LNA2 binding to Bvht H9:cs
Table S5. Detailed fitting results for bootstrapped data sets of LNA2 binding to Bvht H9:cs
Table S6. Detailed fitting results for LNA2 binding to Bvht H9 at varying salt concentration
Table S7. Detailed fitting results for LNA2 binding to Bvht H9:cs at varying salt concentration
Table S8. Detailed fitting results for MO binding to Bvht H9 at varying salt concentration
Table S9. Detailed fitting results for MO binding to Bvht H9:cs at varying salt concentration

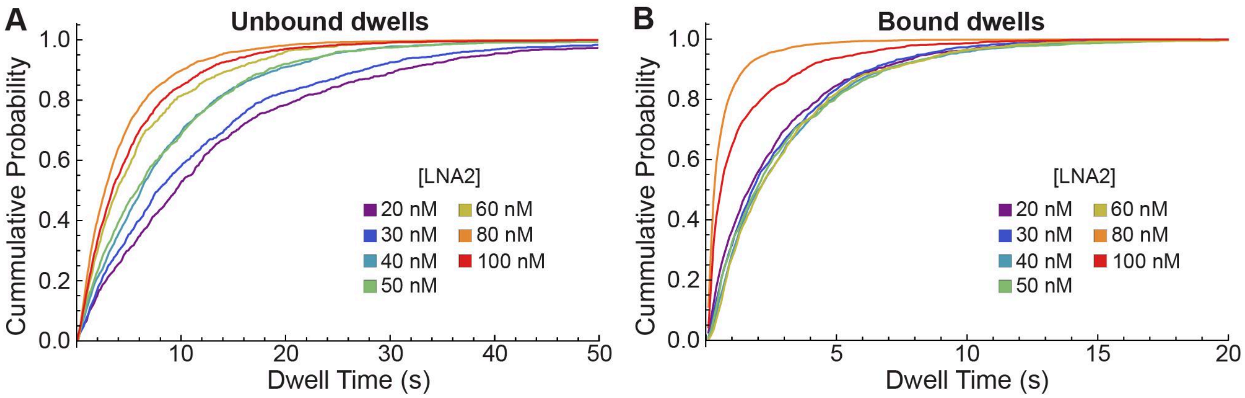


**Figure S1.** Impact of probe concentration on dwell times of LNA2 binding to Bvht H9:cs. *A*, Cumulative distributions of unbound dwell times at indicated LNA2 concentrations. *B*, Bound dwell times at indicated LNA2 concentrations. As expected, unbound dwell time decreases with increasing probe concentration while bound dwell time remains unchanged up to 60 nM probe. The drastic increase in short bound dwells at 80 and 100 nM LNA2 indicates that fluctuations in background noise due to the fluorescence of freely diffusing probe are being identified as binding events. Hence, 60 nM was chosen as a concentration that yielded a substantial number of binding events per trace (quantified in main text tables) without introducing excessive noise.

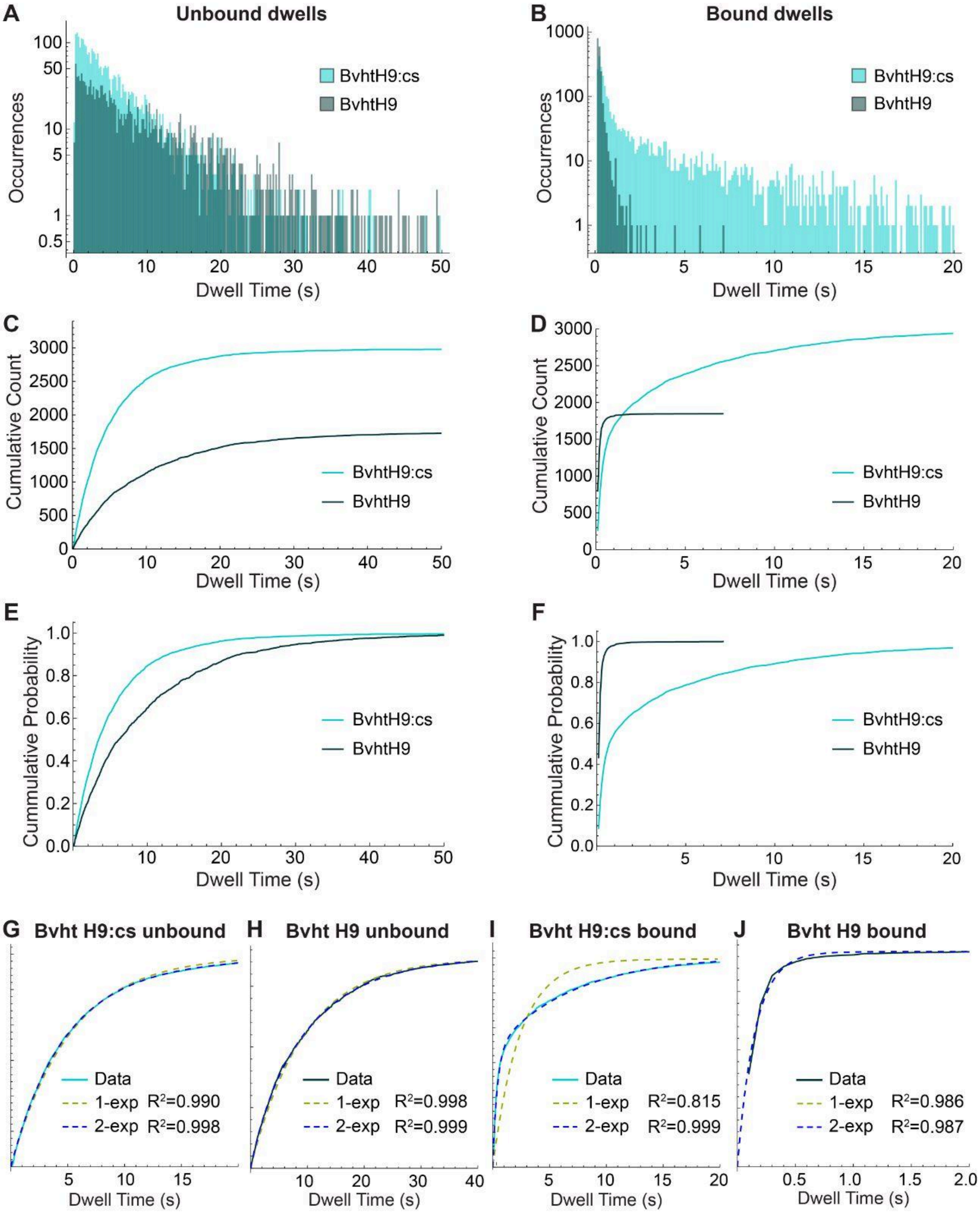


**Fig S2.** Overview of dwell time distribution analysis. Data on LNA2 in the presence of 1 mM $Mg^{2+}$ is used as an example. *A*, Histogram displaying the number of times an unbound segment of a trace lasted for the corresponding dwell time indicated on the x-axis. For clearer visualization, dwell times are binned into 2-frame (0.2 s) increments. Includes all “bracketed” unbound segments of all traces recorded under this experimental condition (segments that are not truncated by the beginning or end of the trace). The y-axis is on a logarithmic scale. *B*, Same as *A* for bound segments of traces. *C*, Cumulative distribution plot displaying the *number* of unbound trace segments with duration less than or equal to the corresponding dwell time indicated on the x-axis. Curves begin at the x-value corresponding to the frame rate of the movie (0.1 s), indicating a dwell time of 1 frame. This plot preserves all of the information present in an unbinned dwell time histogram. *D*, Same as *C* but for bound segments of traces. Curves end at the x-value corresponding to the longest observed bound dwell time. *E*, Cumulative distribution plot displaying the *probability* that an unbound segment of a trace has a duration less than or equal to the corresponding time indicated on the x-axis. This plot preserves all of the information in the histogram except for the total number of dwells that were analyzed. *F*, Same as *E* but for bound segments of traces. *G*-*J*, Exponential fitting of cumulative distribution functions, showing cumulative count curves (solid) and single- (dashed yellow) and double- (dashed blue) exponential fits. The $R^2$ value for each fit is provided. *G*, Bvht H9:cs unbound dwells. *H*, Bvht H9 unbound dwells. *I*, Bvht H9:cs bound dwells. *J*, Bvht H9 bound dwells.

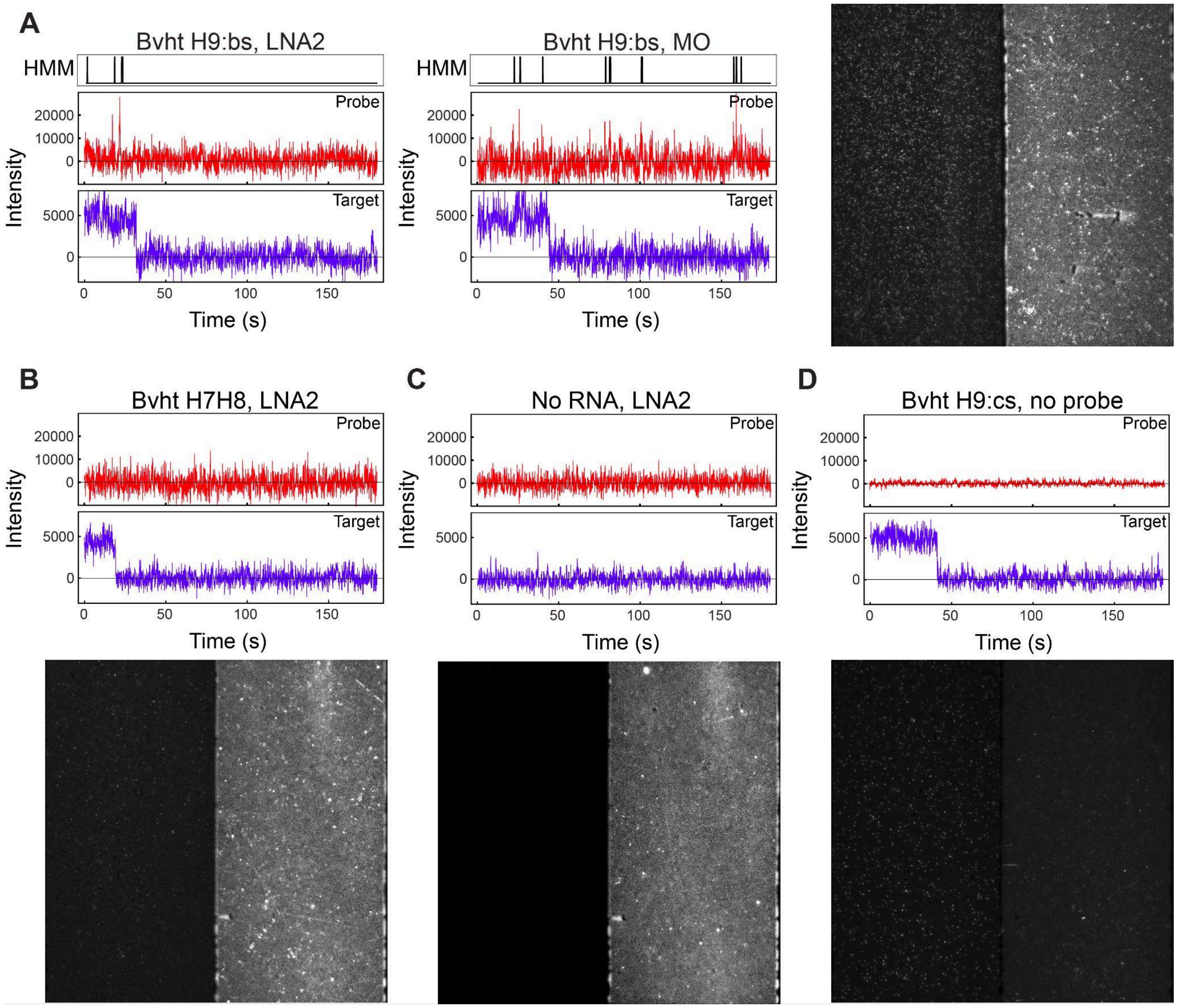


**Figure S3.** Representative single-molecule traces of control conditions. *A*, Representative single-molecule traces for LNA2 (left) or MO (middle) probing Bvht H9:bs. Top: HMM idealization of red (probe) channel trace. Middle: Red (probe) channel trace; Bottom: Green (target) channel trace; Right: example image from SiM-KARTS data collection of LNA2 probing Bvht H9:bs.*B,* Same as *A* but for LNA2 probing Bvht H7H8, with example image below. (N=63 traces analyzed; D/N=2 average bound dwells per trace). *C,* Same as *B* but for LNA2 with no target RNA present (N=167; D/N=1). *D,* Same as *B* but for Bvht H9:cs in the absence of probe (N=124; D/N=1).

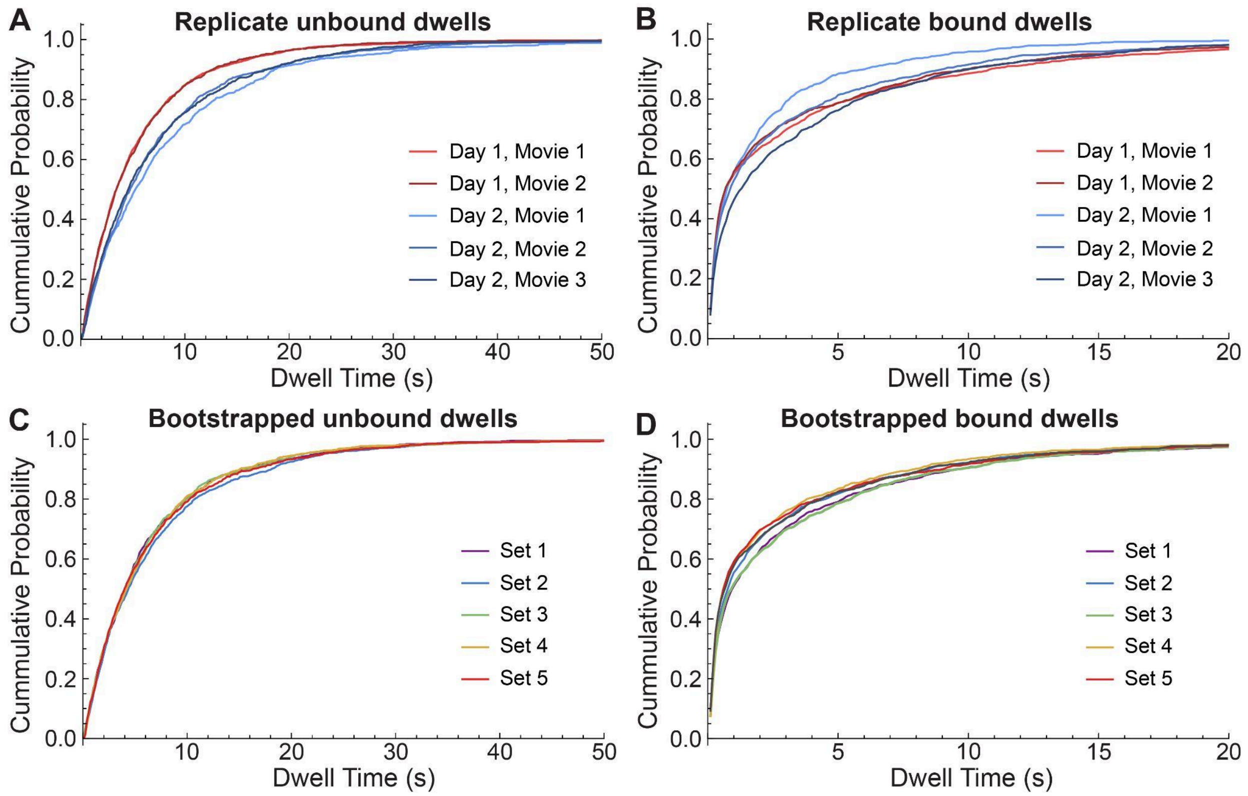


**Figure S4.** SiM-KARTS data reproducibility. *A*, Cumulative distributions of unbound dwell times for LNA2 binding to Bvht H9:cs at 1 mM $Mg^{2+}$ across replicate movies (variations in shade) recorded on two days (red vs blue). *B*, Bound dwell times for LNA2 binding to Bvht H9:cs at 1 mM $Mg^{2+}$ across replicate movies recorded on two days. *C,* Data from *A* after bootstrapping analysis performed by randomly dividing all traces included in panels *A*-*B* into five sets and analyzing each independently. *D*, Data from *B* after bootstrapping analysis.

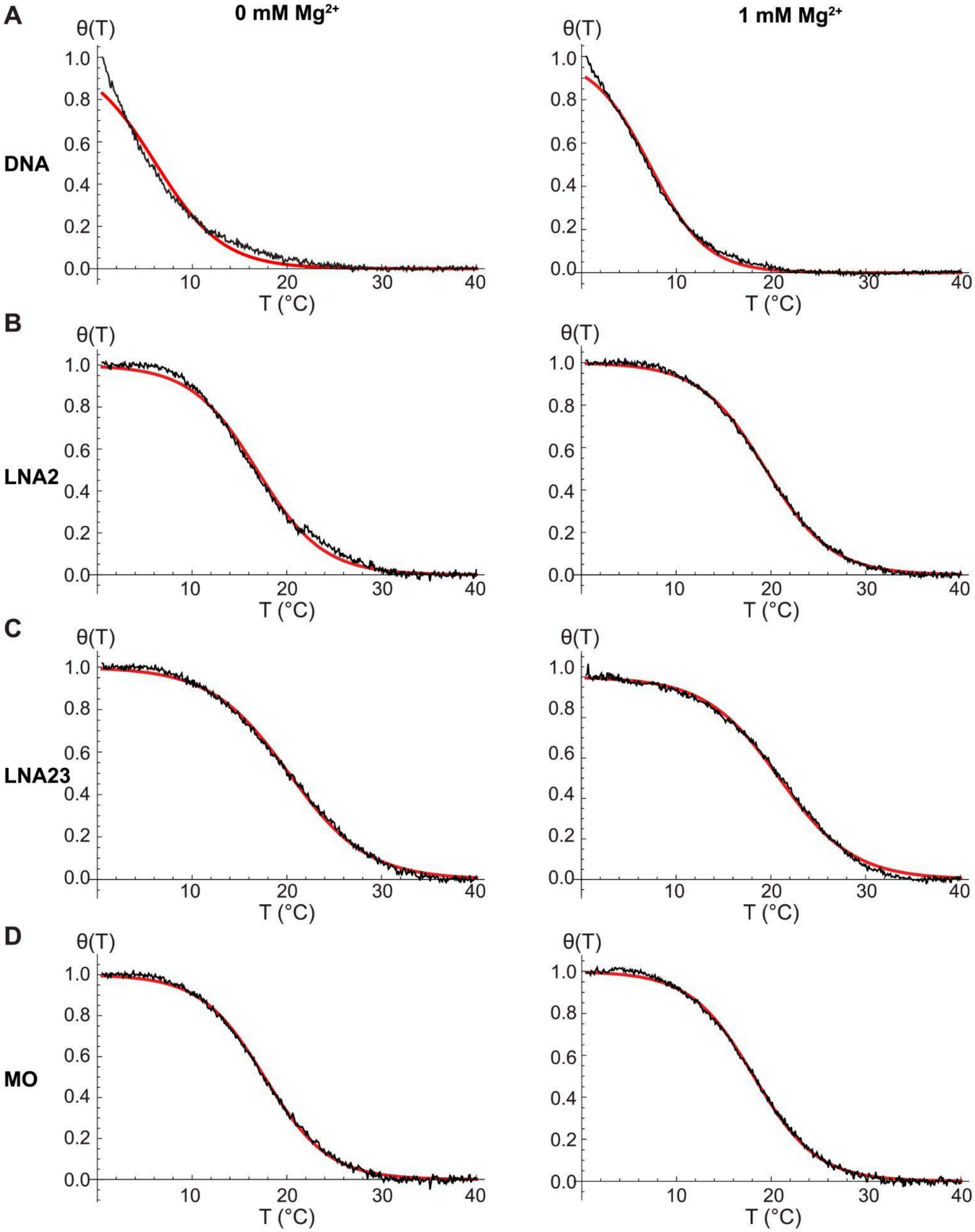


**Figure S5.** Processed melting curves represented as fraction bound θ vs. temperature T (black) and fits (red) recorded at 0 mM $MgCl_2$ (left column) or 1 mM $MgCl_2$ (right column). *A*, DNA probe. *B*, LNA2. *C*, LNA23. *D*, MO.

| Probe | $[Mg^{2+}]$ (mM) | $\tau_{unbound}$ (s) | $R^2_{unbound}$ | $\tau_{bound,1}$ (s) | $w_{bound,1}$ | $\tau_{bound,2}$ (s) | $w_{bound,2}$ | $R^2_{bound}$ |
|---|---|---|---|---|---|---|---|---|
| DNA | 0 | 9.0 | 0.999 | 0.146 | 1.00 | - | - | 0.988 |
| DNA | 1 | 8.9 | 0.994 | 0.170 | 1.00 | - | - | 0.978 |
| LNA2 | 0 | 9.9 | 0.998 | 0.14 | 1.00 | - | - | 0.989 |
| LNA2 | 1 | 9.4 | 0.998 | 0.15 | 1.00 | - | - | 0.987 |
| LNA23 | 0 | 5.8 | 0.997 | 0.213 | 1.00 | - | - | 0.978 |
| LNA23 | 1 | 5.4 | 0.996 | 0.222 | 1.00 | - | - | 0.977 |
| MO | 0 | 7.6 | 0.997 | 1.22 | 0.13 | 0.30 | 0.87 | 0.997 |
| MO | 1 | 9.1 | 0.992 | 2.92 | 0.14 | 0.26 | 0.86 | 0.997 |

**Table S1**. Detailed fitting results for the Bvht H9 data reported in Table 2. $\tau_{unbound}$ is obtained from a single-exponential fit to the cumulative distribution of unbound dwell times. $\tau_{bound,1}$, $\tau_{bound,2}$, $w_1$, and $w_2$ are the time constants and associated weights obtained from a double-exponential fit to the cumulative distribution of bound dwell times. Goodness of fit is presented as $R^2$.

| Probe | $[Mg^{2+}]$ (mM) | $\tau_{unbound}$ (s) | $R^2_{unbound}$ | $\tau_{bound,1}$ (s) | $w_{bound,1}$ | $\tau_{bound,2}$ (s) | $w_{bound,2}$ | $R^2_{bound}$ |
|---|---|---|---|---|---|---|---|---|
| DNA | 0 | 9.5 | 0.999 | 0.69 | 0.19 | 0.21 | 0.81 | 0.996 |
| DNA | 1 | 12 | 0.993 | 0.96 | 0.12 | 0.25 | 0.88 | 0.996 |
| LNA2 | 0 | 3.7 | 0.992 | 2.60 | 0.45 | 0.39 | 0.55 | 0.998 |
| LNA2 | 1 | 5.2 | 0.998 | 6.93 | 0.45 | 0.41 | 0.55 | 0.999 |
| LNA23 | 0 | 3.9 | 0.972 | 7.2 | 0.62 | 0.6 | 0.38 | 0.998 |
| LNA23 | 1 | 8.0 | 0.996 | 16.1 | 0.49 | 1.3 | 0.51 | 0.999 |
| MO | 0 | 5.4 | 0.995 | 2.96 | 0.20 | 0.42 | 0.80 | 0.995 |
| MO | 1 | 7.6 | 0.997 | 2.39 | 0.15 | 0.26 | 0.85 | 0.992 |

**Table S2.** Detailed fitting results for the Bvht H9:cs data reported in Table 2. $\tau_{unbound}$ is obtained from a single-exponential fit to the cumulative distribution of unbound dwell times. $\tau_{bound,1}$, $\tau_{bound,2}$, $w_1$, and $w_2$ are the time constants and associated weights obtained from a double-exponential fit to the cumulative distribution of bound dwell times. Goodness of fit is presented as $R^2$.

| Probe | $[Mg^{2+}]$ (mM) | $\tau_{unbound}$ (s) | $R^2_{unbound}$ | $\tau_{bound,1}$ (s) | $w_{bound,1}$ | $\tau_{bound,2}$ (s) | $w_{bound,2}$ | $R^2_{bound}$ |
|---|---|---|---|---|---|---|---|---|
| LNA2 | 0 | 19.6 | 0.980 | 2.11 | 0.11 | 0.26 | 0.89 | 0.992 |
| LNA2 | 1 | 21.5 | 0.982 | 1.21 | 0.18 | 0.28 | 0.82 | 0.994 |
| MO | 0 | 18.5 | 0.963 | 1.66 | 0.18 | 0.29 | 0.82 | 0.993 |
| MO | 1 | 17.8 | 0.983 | 2.77 | 0.13 | 0.26 | 0.87 | 0.992 |

**Table S3.** Detailed fitting results for the Bvht H9:bs data reported in Table 3. $\tau_{unbound}$ is obtained from a single-exponential fit to the cumulative distribution of unbound dwell times. $\tau_{bound,1}$, $\tau_{bound,2}$, $w_1$, and $w_2$ are the time constants and associated weights obtained from a double-exponential fit to the cumulative distribution of bound dwell times. Goodness of fit is presented as $R^2$.

| Day | Movie | $\tau_{unbound}$ (s) | $R^2_{unbound}$ | $\tau_{bound,1}$ (s) | $w_{bound,1}$ | $\tau_{bound,2}$ (s) | $w_{bound,2}$ | $R^2_{bound}$ | N | D/N |
|---|---|---|---|---|---|---|---|---|---|---|
| 1 | 1 | 5.19 | 0.998 | 7.02 | 0.46 | 0.41 | 0.54 | 0.998 | 95 | 18 |
| 1 | 2 | 5.16 | 0.999 | 6.68 | 0.44 | 0.40 | 0.56 | 0.998 | 71 | 19 |
| 1[1] | 1,2 | 5.20 | 0.998 | 6.93 | 0.45 | 0.41 | 0.55 | 0.999 | 166 | 18 |
| 2 | 1 | 6.97 | 0.998 | 5.51 | 0.48 | 0.44 | 0.52 | 0.999 | 71 | 15 |
| 2 | 2 | 7.71 | 0.998 | 3.64 | 0.50 | 0.44 | 0.50 | 0.997 | 63 | 15 |
| 2 | 3 | 6.91 | 0.998 | 5.71 | 0.58 | 0.43 | 0.42 | 0.999 | 73 | 15 |
| 2 | 1,2,3 | 7.23 | 0.998 | 5.13 | 0.50 | 0.46 | 0.50 | 0.999 | 207 | 15 |

**Table S4.** Detailed fitting results for experimental replicates of LNA2 binding to Bvht H9:cs at 1 mM $Mg^{2+}$. $\tau_{unbound}$ is obtained from a single-exponential fit to the cumulative distribution of unbound dwell times. $\tau_{bound,1}$, $\tau_{bound,2}$, $w_1$, and $w_2$ are the time constants and associated weights obtained from a double-exponential fit to the cumulative distribution of bound dwell times. Goodness of fit is presented as $R^2$. "N" indicates the number of single-molecule traces included in each condition. "D/N" indicates the average number of bound dwells observed per trace.
[1]Data in this row reproduced from Table S2.

| Set | $\tau_{unbound}$ (s) | $R^2_{unbound}$ | $\tau_{bound,1}$ (s) | $w_{bound,1}$ | $\tau_{bound,2}$ (s) | $w_{bound,2}$ | $R^2_{bound}$ | N | D/N |
|---|---|---|---|---|---|---|---|---|---|
| 1 | 6.05 | 0.997 | 6.73 | 0.47 | 0.40 | 0.53 | 0.998 | 74 | 13 |
| 2 | 5.99 | 0.996 | 6.29 | 0.48 | 0.40 | 0.52 | 0.999 | 74 | 14 |
| 3 | 6.03 | 0.997 | 5.58 | 0.45 | 0.37 | 0.55 | 0.999 | 75 | 14 |
| 4 | 5.94 | 0.999 | 5.40 | 0.50 | 0.41 | 0.50 | 0.997 | 75 | 14 |
| 5 | 5.55 | 0.996 | 6.47 | 0.49 | 0.34 | 0.51 | 0.998 | 75 | 13 |

**Table S5.** Detailed fitting results for bootstrapped data sets of LNA2 binding to Bvht H9:cs at 1 mM $Mg^{2+}$. $\tau_{unbound}$ is obtained from a single-exponential fit to the cumulative distribution of unbound dwell times. $\tau_{bound,1}$, $\tau_{bound,2}$, $w_1$, and $w_2$ are the time constants and associated weights obtained from a double-exponential fit to the cumulative distribution of bound dwell times. Goodness of fit is presented as $R^2$. "N" indicates the number of single-molecule traces included in each condition. "D/N" indicates the average number of bound dwells observed per trace.

| $[Mg^{2+}]$ (mM) | $[Na^+]$ (mM) | $\tau_{unbound}$ (s) | $R^2_{unbound}$ | $\tau_{bound,1}$ (s) | $w_{bound,1}$ | $\tau_{bound,2}$ (s) | $w_{bound,2}$ | $R^2_{bound}$ |
|---|---|---|---|---|---|---|---|---|
| 0[1] | 100 | 9.9 | 0.998 | 0.14 | 1.00 | - | - | 0.989 |
| 1[1] | 100 | 9.4 | 0.998 | 0.15 | 1.00 | - | - | 0.987 |
| 1 | 200 | 9.6 | 0.999 | 0.19 | 1.00 | - | - | 0.990 |
| 1 | 300 | 9.3 | 0.993 | 0.21 | 1.00 | - | - | 0.989 |
| 1 | 500 | 10.0 | 0.993 | 0.21 | 0.94 | 1.23 | 0.06 | 0.995 |
| 20 | 100 | 9.1 | 0.993 | 0.17 | 1.00 | - | - | 0.988 |

**Table S6.** Detailed fitting results for the Bvht H9 data reported in Table 7 (LNA2 salt series). $\tau_{unbound}$ is obtained from a single-exponential fit to the cumulative distribution of unbound dwell times. $\tau_{bound,1}$, $\tau_{bound,2}$, $w_1$, and $w_2$ are obtained from a double-exponential fit to the cumulative distribution of bound dwell times. Goodness of fit is presented as $R^2$.
[1]Data in these rows reproduced from Table S1.

| $[Mg^{2+}]$ (mM) | $[Na^+]$ (mM) | $\tau_{unbound}$ (s) | $R^2_{unbound}$ | $\tau_{bound,1}$ (s) | $w_{bound,1}$ | $\tau_{bound,2}$ (s) | $w_{bound,2}$ | $R^2_{bound}$ |
|---|---|---|---|---|---|---|---|---|
| 0[1] | 100 | 3.7 | 0.992 | 2.60 | 0.45 | 0.39 | 0.55 | 0.998 |
| 1[1] | 100 | 5.2 | 0.998 | 6.93 | 0.45 | 0.41 | 0.55 | 0.999 |
| 1 | 200 | 6.8 | 0.999 | 6.6 | 0.51 | 0.5 | 0.49 | 0.999 |
| 1 | 300 | 5.0 | 0.999 | 6.12 | 0.54 | 0.44 | 0.46 | 0.999 |
| 1 | 500 | 5.7 | 0.996 | 7.7 | 0.48 | 0.5 | 0.52 | 0.998 |
| 20 | 100 | 7.4 | 0.971 | 0.47 | 0.11 | 0.10 | 0.89 | 1.000 |

**Table S7.** Detailed fitting results for the Bvht H9:cs data reported in Table 7 (LNA2 salt series). $\tau_{unbound}$ is obtained from a single-exponential fit to the cumulative distribution of unbound dwell times. $\tau_{bound,1}$, $\tau_{bound,2}$, $w_1$, and $w_2$ are obtained from a double-exponential fit to the cumulative distribution of bound dwell times. Goodness of fit is presented as $R^2$.
[1]Data in these rows reproduced from Table S2.

| $[Mg^{2+}]$ (mM) | $[Na^+]$ (mM) | $\tau_{unbound}$ (s) | $R^2_{unbound}$ | $\tau_{bound,1}$ (s) | $w_{bound,1}$ | $\tau_{bound,2}$ (s) | $w_{bound,2}$ | $R^2_{bound}$ |
|---|---|---|---|---|---|---|---|---|
| 0[1] | 100 | 7.6 | 0.997 | 1.22 | 0.13 | 0.30 | 0.87 | 0.997 |
| 1[1] | 100 | 9.0 | 0.992 | 2.92 | 0.14 | 0.26 | 0.86 | 0.997 |
| 1 | 500 | 7.7 | 0.991 | 2.65 | 0.12 | 0.33 | 0.88 | 0.990 |

**Table S8.** Detailed fitting results for the Bvht H9 data reported in Table 8 (MO salt series). $\tau_{unbound}$ is obtained from a single-exponential fit to the cumulative distribution of unbound dwell times. $\tau_{bound,1}$, $\tau_{bound,2}$, $w_1$, and $w_2$ are obtained from a double-exponential fit to the cumulative distribution of bound dwell times. Goodness of fit is presented as $R^2$.
[1]Data in these rows reproduced from Table S1.

| $[Mg^{2+}]$ (mM) | $[Na^+]$ (mM) | $\tau_{unbound}$ (s) | $R^2_{unbound}$ | $\tau_{bound,1}$ (s) | $w_{bound,1}$ | $\tau_{bound,2}$ (s) | $w_{bound,2}$ | $R^2_{bound}$ |
|---|---|---|---|---|---|---|---|---|
| 0[1] | 100 | 5.4 | 0.995 | 2.96 | 0.20 | 0.42 | 0.80 | 0.995 |
| 1[1] | 100 | 7.6 | 0.997 | 2.39 | 0.15 | 0.26 | 0.85 | 0.992 |
| 1 | 500 | 7.5 | 0.998 | 2.82 | 0.11 | 0.27 | 0.89 | 0.991 |

**Table S9.** Detailed fitting results for the Bvht H9:cs data reported in Table 8 (MO salt series). $\tau_{unbound}$ is obtained from a single-exponential fit to the cumulative distribution of unbound dwell times. $\tau_{bound,1}$, $\tau_{bound,2}$, $w_1$, and $w_2$ are obtained from a double-exponential fit to the cumulative distribution of bound dwell times. Goodness of fit to the respective cumulative distribution is presented as $R^2$.

[1]Data in these rows reproduced from Table S2.
